# The segregase CDC48 integrates blue light and hormonal cues to regulate photomorphogenesis in Arabidopsis

**DOI:** 10.64898/2026.04.23.720413

**Authors:** Antonela L. Alem, Agustín L. Arce, Cécile Blanchard, María Dolores Gómez, Esther Carrera, Olivier Lamotte, Miguel A. Pérez-Amador, Matías Capella

## Abstract

Photomorphogenesis allows plants to adjust growth to ambient light conditions and relies on protein quality control to ensure the timely turnover of signaling components. The conserved AAA+ ATPase CDC48, along with its cofactors NPL4 and UFD1, is a crucial regulator of proteasomal degradation. While well characterized in other organisms, its role in plant development remains largely unexplored. Here, we show that CDC48 is required for blue light-mediated photomorphogenesis in Arabidopsis. Under blue light, CDC48A accumulates at the plasma membrane and in the nucleus, and *cdc48a* mutants fail to repress hypocotyl elongation properly. Similar phenotypes are observed upon inhibition of CDC48 or in *npl4* and *ufd1* mutants. Genetic and biochemical analyses further reveal that CDC48A negatively regulates gibberellin (GA) signaling. Consistently, UFD1 directly interacts with the GA receptor GID1 to promote its degradation. Together, these findings demonstrate that CDC48A integrates light and hormonal cues through protein homeostasis to regulate photomorphogenic development.

## Introduction

Besides its fundamental role as an energy source, light functions as a key environmental signal that shapes plant architecture. The spectral composition, irradiance, and directionality of light profoundly influence every stage of growth and development. In the absence of light, germinating seedlings undergo skotomorphogenesis, a developmental program characterized by rapid hypocotyl elongation, the maintenance of an apical hook, and closed cotyledons. Upon light perception, seedlings transition to photomorphogenesis, which triggers hypocotyl growth inhibition, cotyledon expansion, and chloroplast biogenesis (Krahmer and Fankhauser 2024). Light is perceived by multiple photoreceptors, including the red/far-red-sensing phytochromes (PHYA-PHYE) and the blue-light-sensing cryptochromes (CRY1-CRY2), which initiate specialized signaling cascades to modulate developmental plasticity. For instance, these photoreceptors modulate the activity of several members of the PHYTOCHROME INTERACTING FACTOR (PIF) family of basic helix-loop-helix (bHLH) transcription factors, which promote hypocotyl elongation through the activation of growth-promoting genes (Leivar et al. 2008b; Shin et al. 2009; Kunihiro et al. 2010).

Photomorphogenic growth is also influenced by the integration of light with endogenous hormonal cues, most notably auxin, brassinosteroids, and gibberellins (GAs) (Arsovski et al. 2012). GAs promote growth by facilitating the binding between its receptor, GA INSENSITIVE DWARF 1 (GID1), and the growth-repressing DELLA proteins, thereby triggering DELLA degradation. Under blue light, CRY1 antagonizes this process by directly interacting with and preventing GID1-DELLA association, resulting in DELLA accumulation (Xu et al. 2021; Yan et al. 2021; Zhong et al. 2021). Once stabilized, DELLAs sequester PIFs, preventing their binding to target promoters or promoting their degradation (Feng et al. 2008; Li et al. 2016). The selective degradation of DELLAs and PIFs is mainly mediated by the ubiquitin-proteosome system (UPS), a key regulatory pathway underlying the extensive proteome reorganization triggered by light perception (Zhou and Deng 2025). Central to this landscape are the COP/DET/FUS complexes, which function as essential repressors of photomorphogenesis in the dark. Specifically, the RING-type E3 ubiquitin ligase CONSTITUTIVE PHOTOMORPHOGENIC1 (COP1) mediates the ubiquitination and subsequent proteasomal degradation of photomorphogenesis-promoting factors, including the central light regulator ELONGATED HYPOCOTYL 5 (HY5) (Ponnu and Hoecker 2021).

The evolutionarily conserved AAA+ ATPase CDC48 (cell division cycle 48; p97/VCP in humans) functions as a molecular segregase central to ubiquitin-dependent protein quality control. Operating as hexamers, CDC48 uses ATP hydrolysis to extract and unfold ubiquitinated or SUMOylated substrates from membranes, chromatin, or multi-protein complexes. Structurally, each monomer comprises two tandem ATPase domains (D1 and D2) that assemble into stacked hexameric rings forming a central translocation channel. Conserved Walker A and Walker B motifs within these domains coordinate the ATP binding and hydrolysis required to generate the mechanical force for protein disassembly (Meyer and van den Boom 2023).

CDC48 orthologs have been extensively studied in yeast and mammals, where they participate in a wide diversity of cellular processes (Meyer and Weihl 2014). For example, CDC48 modulates chromatin dynamics during DNA damage or transcriptional activation (Bonizec et al. 2014; Mérai et al. 2014; Capella et al. 2021). In plants, CDC48 has been shown to contribute to immune responses, as well as protein degradation pathways associated with chloroplasts and mitochondria, the endoplasmic reticulum, and the inner nuclear membrane (Müller et al. 2005; Niehl et al. 2012; Copeland et al. 2016; Ling et al. 2019; Huang et al. 2020; Ao et al. 2021; Li et al. 2022; Li and Jarvis 2024; Schoberer et al. 2024; Yang et al. 2024; Blanchard et al. 2025). This functional versatility is achieved through the recruitment of specialized cofactors, such as the UFD1-NPL4 complex, which guides substrate recognition and processing. For instance, the plant CDC48-UFD1-NPL4 complex (CDC48-UN) has been implicated in heterochromatin decondensation to activate ribosomal DNA and the degradation of proteins located in chloroplasts or the nuclear membrane (Mérai et al. 2014; Huang et al. 2020; Li et al. 2022).

In Arabidopsis, the CDC48 family comprises three homologues (CDC48A-C), with CDC48A being the primary isoform regulating cell division and growth (Rancour et al. 2002; Park et al. 2008). Beyond its role in basal cellular maintenance, whether CDC48 contributes to specific developmental programs remains unknown. Here, we demonstrate that CDC48 is a critical regulator of the photomorphogenic response to blue light. We show that blue light triggers CDC48 enrichment at the nuclear and plasma membrane, where its segregase activity is required to suppress hypocotyl growth. Furthermore, our data indicate that CDC48-UN negatively regulates GA signaling by mediating the degradation of the GA receptor GID1. Together, our findings identify CDC48 as a new player in the control of plant development, by modulating the GA pathway to facilitate blue-light-dependent photomorphogenesis.

## Results

### CDC48A exhibits a blue light-specific subcellular localization pattern in hypocotyl cells

Photomorphogenesis is a light-driven developmental program that relies on protein homeostasis to ensure proper folding, activation, and timely degradation of key signaling components (Zhou and Deng 2025). Given the established role of CDC48 in protein quality control, we postulated that CDC48A-mediated proteostasis may also regulate plant developmental processes. To address this, we focused on light-regulated growth, where protein turnover is known to be an important regulatory mechanism. The localization of CDC48 is known to be highly dynamic, shifting between compartments to facilitate targeted protein quality control across a wide array of plant biological processes (Park et al. 2008; Niehl et al. 2012; Inès et al. 2025). Therefore, to determine whether light influences CDC48A spatial distribution, we explored its subcellular localization in Arabidopsis seedlings expressing *^YFP^CDC48A* under its native promoter grown in either complete darkness or under long-day conditions. We found that light promoted a distinct localization pattern of ^YFP^CDC48A in hypocotyl cells compared with darkness, characterized by predominant accumulation at the plasma membrane and within the nucleus (Fig. 1a). This effect was wavelength-specific, as only blue light induced the characteristic localization (Fig. 1a). Similar results were obtained in *Nicotiana benthamiana* leaves transiently overexpressing *^GFP^CDC48A* (Fig. 1b). Tobacco co-transformation with the nuclear marker NLS-mCherry corroborated the light-induced nuclear enrichment of CDC48A (Supplementary Fig. S1a). Despite the clear requirement for blue light to trigger this redistribution, the subcellular localization of ^YFP^CDC48A was unaffected in *cry1-304* seedlings (Supplementary Fig. S1b). These results reveal that CDC48 undergoes a specific spatial redistribution in response to blue-light perception.

**Fig. 1.**
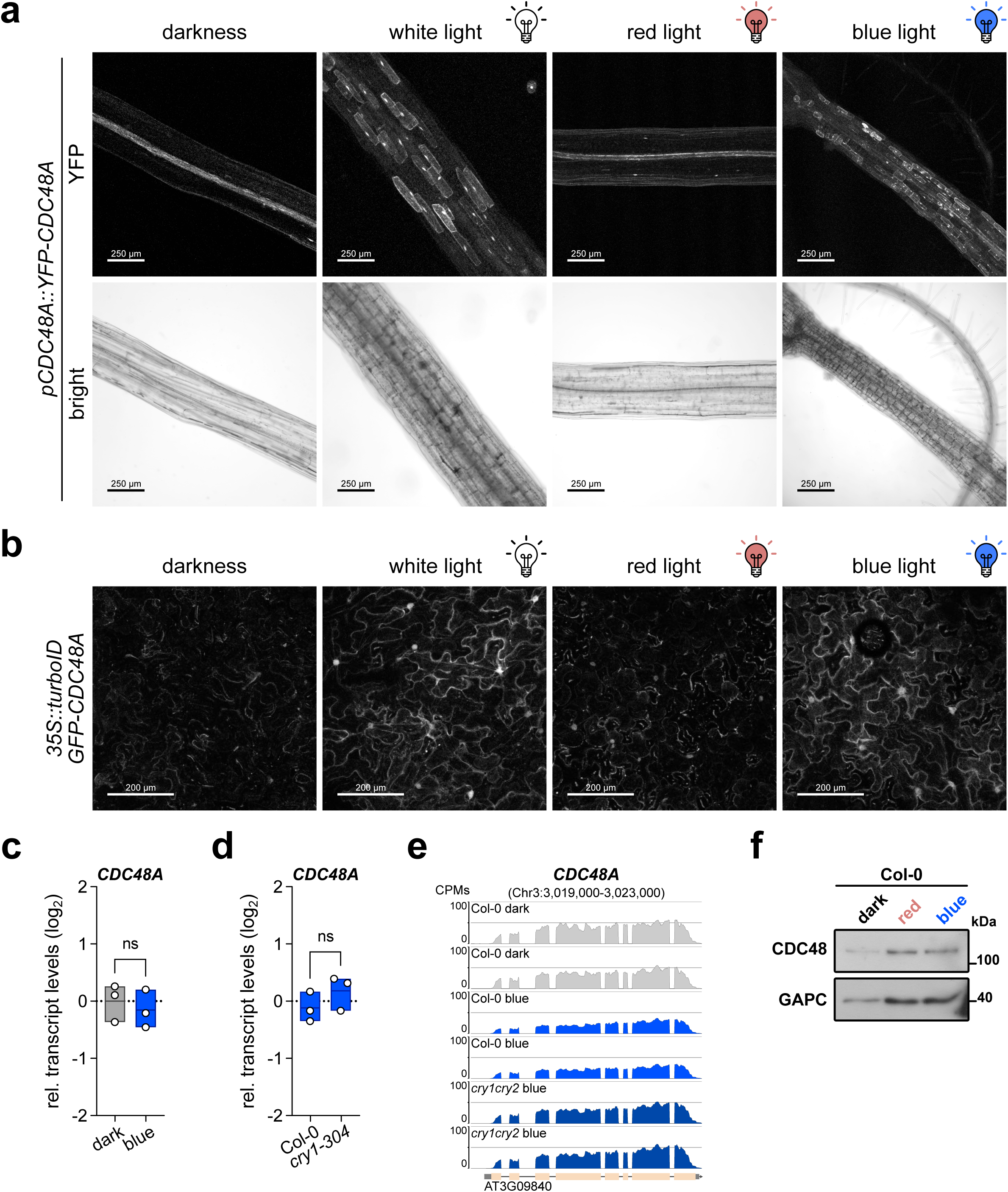
Blue light promotes CDC48A accumulation at the nucleus and plasma membrane in plants. **a**, Representative images of the live-cell imaging of 4-day-old *pCDC48A::YFP-CDC48A* seedlings grown under continuous darkness, white, red, or blue light. Scale bars, 250 µm. **b**, Representative images of tobacco leaves transformed with *35S::turboID-GFP-CDC48A* grown under continuous darkness, white, red, or blue light. Scale bars, 200 µm. **c-d**, *CDC48A* transcript levels quantified by RT-qPCR in Col-0 wild-type seedlings grown in darkness or blue light (**c**) or Col-0 and *cry1-304* seedlings grown under blue light for 4 d. Data were normalized to *ACTIN2/8* transcript levels and are shown on a log_2_ scale relative to Col-0 grown in darkness, which was set to zero. Data are presented as floating bar plots, with the line indicating the mean of *n* = 3 independent biological replicates and individual values shown. Statistical analysis was performed using a two-tailed Student’s *t*-test. ns, denotes no significant differences. **e**, RNA-seq coverage plots showing transcript levels of *CDC48A* in Col-0 wild-type grown under complete darkness, or Col-0 and *cry1 cry2* seedlings grown under blue light for 6 d at 22 °C. Data were reanalysed from a publicly available dataset deposited in the Gene Expression Omnibus (GEO) under the accession number GSE226927. All reads are presented as counts per million (CPM), and genomic coordinates are shown in base pairs (bp). **f**, Immunoblot of CDC48 from Col-0 seedlings grown under darkness, red, or blue light. GAPC served as loading control. For **a** and **b**, sum-intensity Z-projections are shown.

The light-induced accumulation of CDC48A could result from increased transcription or enhanced protein stability. We measured *CDC48A* transcript levels in 4-day-old wild-type (Col-0) seedlings grown under continuous darkness or blue light and found that *CDC48A* expression remains stable regardless of illumination conditions (Fig. 1c). Additionally, *CDC48A* transcript levels were unaffected in the *cry1-304* mutant under blue light (Fig. 1d), which was corroborated by a re-analysis of public RNA-Seq datasets from WT and *cry1 cry2* seedlings (Fig. 1e). Similarly, red light treatment did not alter transcript abundance (Supplementary Fig. S1c), indicating that *CDC48A* transcription is not light-regulated. Consistent with the transcript data, immunoblot analysis using anti-Cdc48 revealed that total protein abundance remained constant across all light qualities compared to etiolated controls (Fig. 1e). Taken together, our data suggest that blue light triggers the subcellular relocalization of CDC48A to the plasma membrane and nucleus without altering its overall protein levels.

### CDC48A represses hypocotyl growth in blue light

During photomorphogenesis, blue light perception limits hypocotyl elongation and promotes cotyledon expansion and chloroplast development (Liscum and Hangarter 1991). Since CDC48A accumulates in hypocotyl cells under blue light, we examined whether its activity is required for hypocotyl elongation. Since CDC48A is an essential gene, we utilized the partial loss-of-function allele *cdc48a-4* (G274E) (Park et al. 2008; Copeland et al. 2016). Under continuous blue light, *cdc48a-4* exhibited longer hypocotyls than WT seedlings, likely due to increased cell elongation (Fig. 2a-c). No differences were observed in darkness, while *cdc48a-4* showed slightly shorter hypocotyls than controls under red light (Fig. 2b). Notably, the long-hypocotyl phenotype of *cdc48a-4* became more pronounced with increasing light fluence rates (Fig. 2d), resembling the *cry1-304* mutant phenotype. Moreover, *^YFP^CDC48A* overexpression complemented the phenotype of *cdc48a-4* plants (Supplementary Fig. S2a,b). Consistent with ATP-dependent activity, overexpression of *CDC48A* variants defective in ATP binding (K254A/K527A) (Ling et al. 2019) or hydrolysis (E581Q) (Müller et al. 2005) also resulted in longer hypocotyls compared to WT (Fig. 2e,f). Moreover, inhibition of CDC48 activity using CB-5083, an inhibitor of the AAA-ATPase p97/VCP (Zhou et al. 2015), promoted hypocotyl elongation in WT seedlings, while *cdc48a-4* displayed a hypersensitive response to low concentrations of the inhibitor (Fig. 2g). These observations prompted us to examine whether other CDC48 family members contribute to the blue light-mediated photomorphogenic response. Interestingly, while *CDC48B* and *CDC48C* appeared redundant (Supplementary Fig. S2c,d), their mutation in a *cdc48a-4* background further exacerbated hypocotyl elongation (Fig. 2h). Altogether, these findings demonstrate that CDC48 activity is required to inhibit hypocotyl growth upon blue light perception, with CDC48A serving as the primary mediator.

**Fig. 2.**
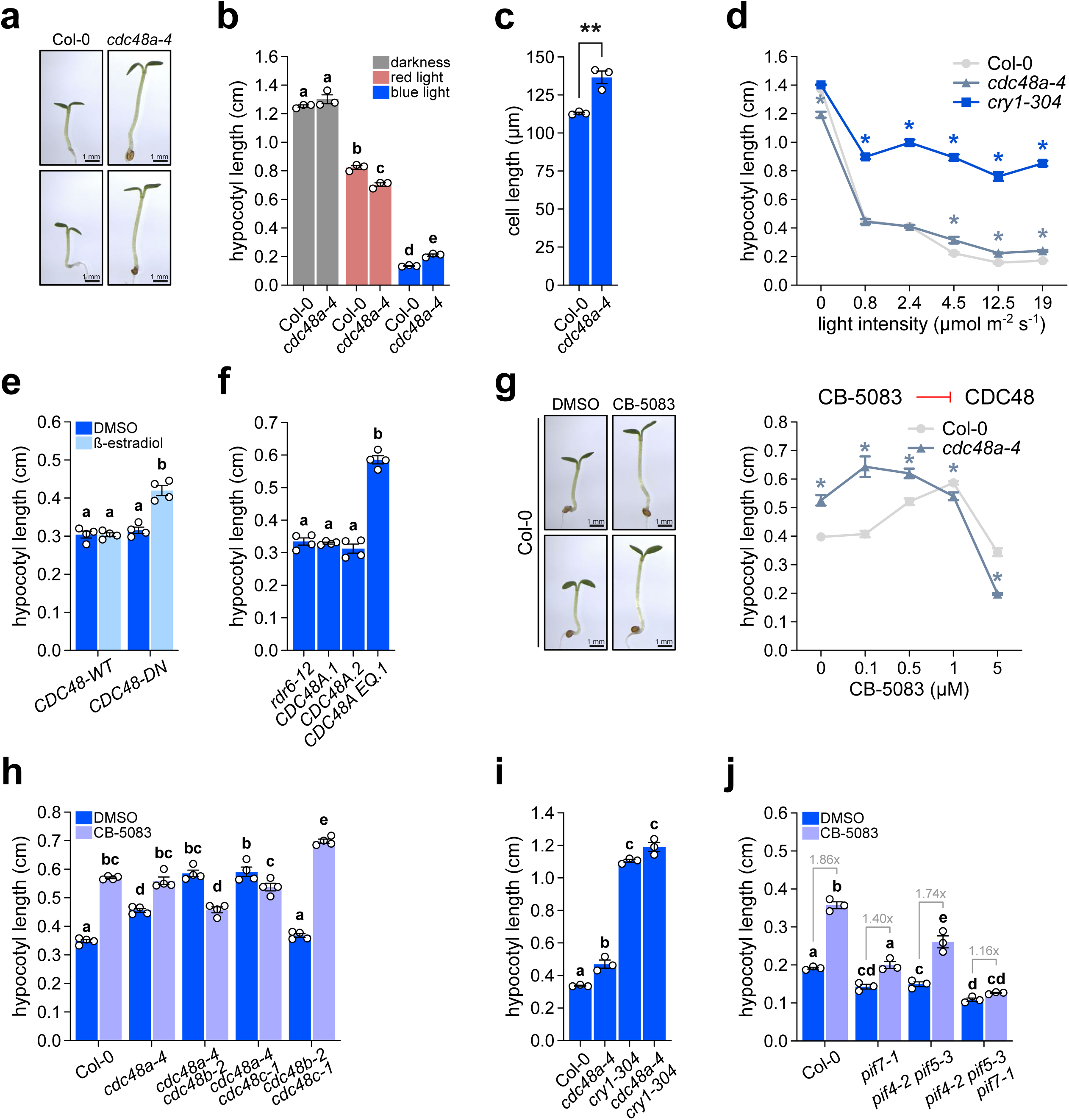
CDC48A activity is required to suppress hypocotyl elongation under blue light conditions. **a**, Representative images of Col-0 and *cdc48a-4* seedlings grown under blue light at 22 °C for 4 d. Scale bar, 1 mm. **b**, Hypocotyl lengths of Col-0 and *cdc48a-4* seedlings grown in darkness, red, or blue light. **c**, Hypocotyl cell length quantification of the indicated seedlings. **d**, Hypocotyl lengths of Col-0, *cdc48a-4*, and *cry1-304* seedlings grown under increasing blue light fluences. **e**, Hypocotyl lengths of seedlings expressing *CDC48-WT* or *CDC48-DN*, induced with β-estradiol (1 µM). **f**, Hypocotyl lengths of *rdr6-12* seedlings untransformed or expressing GFP fusions of *CDC48A* or the E581Q variant (*CDC48A EQ*). **g**, Seedlings grown with the indicated concentrations of the CDC48 inhibitor CB-5083. DMSO (0.05 % v/v) treatment served as a control. Left, representative Col-0 seedlings treated with CB-5083 (1 µM). Scale bar, 1 mm. Right, hypocotyl length quantification in Col-0 and *cdc48a-4*. **h**, Hypocotyl lengths of Col-0 and the indicated *cdc48* mutants grown in the presence or absence of CB-5083 (1 µM). **i**, Hypocotyl lengths of Col-0, *cdc48a-4*, *cry1-304*, and *cdc48a-4 cry1-304*. **j**, Hypocotyl lengths of Col-0 and the indicated *pif* mutants grown as in **h**. Gray numbers indicate fold changes. For **c**, **e-f**, **g-i**, seedlings were grown as in **a**. All data are means (±SEM) of *n* = 3-4 independent biological replicates. For **c**, **d**, and **g**, statistical analysis was performed using a two-tailed Student’s *t*-test. Different letters indicate significant differences among means as determined using one-way ANOVA followed by Tukey’s *post-hoc* test (*P*<0.05).

The observation that CDC48A-mediated suppression of hypocotyl elongation occurs predominantly under medium-to-high blue light intensities suggests that CDC48 may function within a CRY1-dependent pathway(Krahmer and Fankhauser 2024). Indeed, we found that neither the *cry1-304 cdc48a-4* double mutant nor CB-5083 treatment exacerbated the *cry1* phenotype (Fig. 2i; Extended Data Fig 3a,b), implying an epistatic interaction. Furthermore, *CRY1*-overexpressing lines were unaffected by CDC48 inhibition, while *CRY2* overexpressors exhibited a response comparable to WT seedlings (Extended Data Fig 3c,d). Since CRY1 promotes photomorphogenesis by inactivating COP1, which otherwise targets for degradation multiple light-responsive transcription factors like HY5 (Ang et al. 1998; Osterlund et al. 2000; Holtkotte et al. 2017), we tested the involvement of these downstream components. Neither *cop1* nor *hy5* mutants responded to the inactivation of CDC48A activity (Supplementary Fig. S3e,f), supporting the notion that CDC48A acts downstream of CRY1. To gain mechanistic insight into CDC48A-mediated growth control, we examined its genetic interaction with PIFs, which are the central hubs of light-regulated development (Cai and Huq 2024). For this end, we measured hypocotyl length of different single and higher-order *pif* mutants, in the presence or absence of CB-5083. As previously reported (Kunihiro et al. 2010; Burko et al. 2022), PIF4/5/7 are required for hypocotyl elongation under blue light (Fig. 2j). Interestingly, *pif4 pif5 pif7* seedlings were largely insensitive to CDC48 inhibition, with *PIF7* loss likely contributing most prominently to the unresponsive phenotype (Fig. 2j; Supplementary Fig. S3g). In contrast, *pif1* and *pif3* exhibited a response to CB-5083 comparable to WT controls (Supplementary Fig. S3g). Together, our findings indicate that CDC48A acts within the canonical blue light-dependent photomorphogenic pathway.

### CDC48A acts through its co-factors NPL4 and UFD1 to modulate hypocotyl elongation in blue light

The ability of CDC48A to engage in distinct pathways depends, in part, on its association with specific adaptor proteins. Because NPL4 and UFD1 are among the principal cofactors of CDC48(Meyer and van den Boom 2023), we examined their potential role during photomorphogenesis. Live-cell imaging of *NPL4B^GFP^* overexpressing seedlings revealed a subcellular distribution nearly identical to that of CDC48A when grown under white or blue light (Fig. 3a). Moreover, NPL4B protein abundance remained stable across different light conditions (Supplementary Fig. S4a). Consistently, UFD1B also exhibited blue light accumulation in transiently transformed tobacco leaves (Fig. 3b). Furthermore, the double mutants of *NPL4* (*npl4ab*) or *UFD1* (*ufd1bc*) failed to properly inhibit hypocotyl elongation under blue light and were insensitive to CB-5083, phenocopying *cdc48a-4* seedlings (Fig. 3c; Supplementary Fig. S4b,c). These findings suggest that CDC48A-UN modulates the hypocotyl growth of blue light-grown seedlings.

**Fig. 3.**
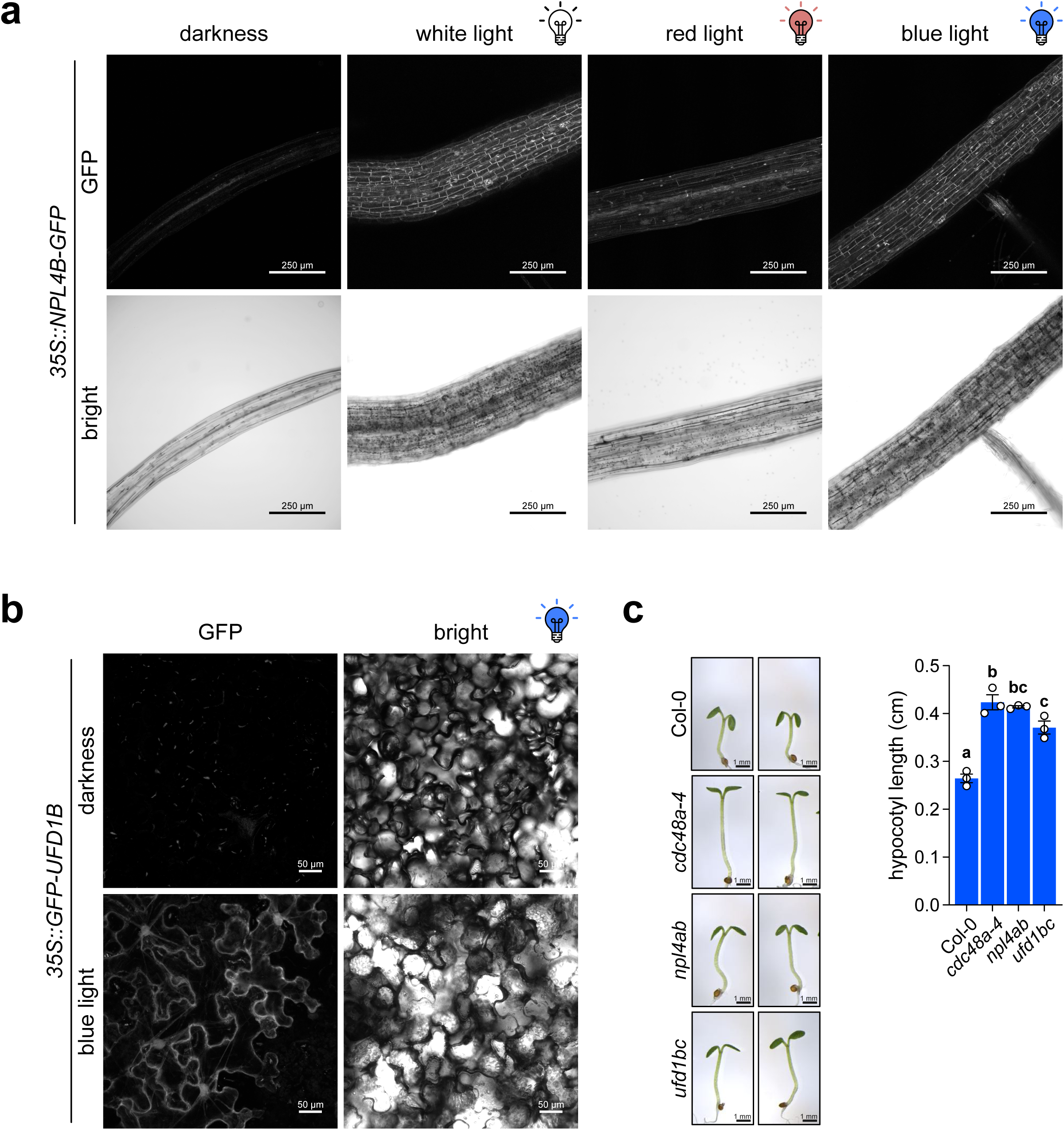
The CDC48A cofactors NPL4 and UFD1 function in blue-light-mediated inhibition of hypocotyl elongation. **a**, Representative images acquired at the end of the live-cell imaging experiment of 4-day-old *35S::NPL4B-GFP* seedlings grown under continuous darkness, white, red, or blue light. Scale bars, 250 µm. **b**, Representative images acquired at the end of the experiment of tobacco leaves transformed with *35S::GFP-UFD1B* grown under continuous darkness or blue light conditions. Scale bars, 50 µm. **c**, Representative images (left) and quantification of hypocotyl length (right) of 4-day-old Col-0 wild-type, *cdc48a-4*, *npl4a npl4b* (*npl4ab*), *ufd1b ufd1c* (*ufd1bc*) seedlings grown under continuous blue light. Data are means (±SEM) of *n* = 3 independent biological replicates, each consisting of at least 15 seedlings grown on the same plate. Different letters indicate significant differences among means as determined using one-way ANOVA followed by Tukey’s *post-hoc* test (*P*<0.05). For **a** and **b**, sum-intensity Z-projections are shown.

### CDC48A negatively regulates the gibberellin hormonal pathway

Auxin is a central promoter of hypocotyl elongation in response to environmental cues, with its biosynthesis and signaling frequently modulated by PIFs (Krahmer and Fankhauser 2024). To evaluate whether CDC48A suppresses hypocotyl growth by modulating auxin homeostasis, we examined the effects of the auxin transport inhibitor 1-naphthylphthalamic acid (NPA) and the synthetic auxin picloram (PIC) on *cdc48a-4* and *npl4ab* seedlings. Both treatments affected hypocotyl elongation to a similar extent in all genotypes tested (Supplementary Fig. S5a,b), supporting the notion that CDC48A functions independently of auxin homeostasis. This was further supported by the observation that the auxin-overproducing *yucca6-1D* mutant exhibited diminished sensitivity to CB-5083, likely reflecting saturation of CDC48-mediated hypocotyl growth (Supplementary Fig. S5c). Moreover, the *AUXIN RESISTANT 1* (*axr1*) mutants, which are deficient in auxin signaling, exhibited hypocotyl elongation comparable to WT upon CDC48 inhibition (Supplementary Fig. S5d). Finally, the activity of the auxin-responsive synthetic *DR5::GUS* reporter remained unchanged following CB-5083 treatment under blue light (Supplementary Fig. S5e). Altogether, these results suggest that CDC48A-mediated hypocotyl elongation inhibition occurs independently of auxin distribution, sensitivity, or signaling.

Besides auxin, gibberellins facilitate hypocotyl elongation in response to developmental and environmental stimuli by promoting GID1-mediated degradation of DELLA proteins(Shani et al. 2024). Therefore, we next investigated whether CDC48A influences GA homeostasis, perception, or signaling to regulate hypocotyl growth under blue light. To test this, we treated *cdc48a-4*, *npl4ab,* and *ufd1bc* mutants with the GA biosynthesis inhibitor paclobutrazol (PAC). Reduction of GA levels impaired the exacerbated hypocotyl growth of all three mutants, restoring a WT-like phenotype (Fig. 4a). Similarly, the CB-5083-induced hypocotyl elongation in WT seedlings was abolished by PAC treatment (Fig. 4b). Consistently, the GA-deficient mutants *ga3ox1-3 ga3ox2-1* and *ga1-3* exhibited a diminished response to CB-5083 treatment compared to WT seedlings (Supplementary Fig. S6a,b). PUX1 (Plant UBX domain-containing protein 1) is a negative regulator of CDC48A that triggers hexamer disassembly (Rancour et al. 2004). Since PUX1 is a repressor of GA signaling and *pux1* mutants display PAC tolerance (Hauvermale et al. 2022), it may act as an endogenous inhibitor of CDC48A-mediated hypocotyl regulation. However, *pux1-3* exhibited WT hypocotyl elongation under blue light (Supplementary Fig. S7). These findings suggest that CDC48A-dependent control of hypocotyl growth requires a functional GA pathway.

**Fig. 4.**
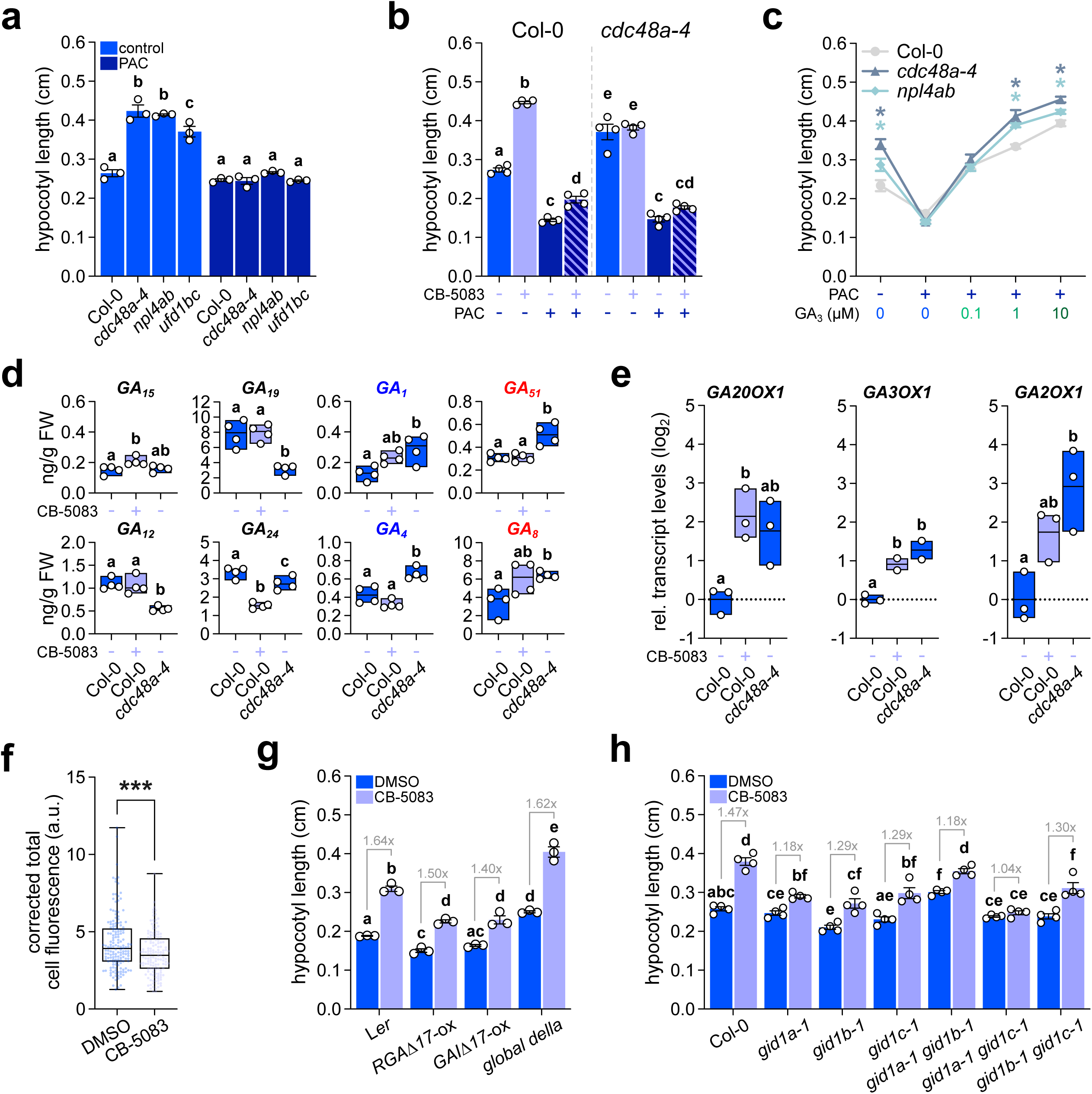
CDC48A negatively regulates hypocotyl elongation in blue light via modulation of gibberellin homeostasis. Seedlings were grown under continuous blue light at 22 °C for 4 d. **a**, Hypocotyl length of Col-0 wild-type, *cdc48a-4*, *npl4a npl4b* (*npl4ab*), and *ufd1b ufd1c* (*ufd1bc*) seedlings grown with or without the GA biosynthesis inhibitor paclobutrazol (PAC; 1 µM). **b**, Hypocotyl length of Col-0 and *cdc48a-4* seedlings grown with or without the CDC48 activity inhibitor CB-5083 (1 µM), PAC (1 µM), or both. **c**, Quantification of hypocotyl length of Col-0, *cdc48a-4*, and *npl4ab* seedlings grown with or without PAC (1 µM) and the indicated concentrations of exogenous GA_3_. **d**, Levels of the indicated GA metabolites in Col-0 and *cdc48a-4* seedlings grown in the presence or absence of CB-5083 (1 µM), expressed as ng GA/g fresh weight. Blue and red labels highlight bioactive GA and catabolites, respectively. **e**, *GA3OX1*, *GA20OX1*, and *GA2OX1* transcript levels quantified by RT-qPCR in Col-0 and *cdc48a-4* seedlings grown as in **d**. Data were normalized to *ACTIN2/8* transcript levels and are shown on a log_2_ scale relative to Col-0 grown under control conditions. **f**, Quantification of GFP signal from live-cell imaging of *pRGA::GFP-RGA* seedlings grown as in **d**. **g**, Hypocotyl length of L*er* wild-type, L*er* seedlings transformed with *35S::TAP-RGA*Δ*17* (*RGA*-ox) or *35S::TAP-GAI*Δ*17* (*GAI*-ox), and the quintuple *della* mutant (*global della*) grown as in **d**. **h**, Hypocotyl lengths of Col-0 and the *gid1a-1*, *gid1b-1*, *gid1c-1*, *gid1a-1 gid1b-1*, *gid1a-1 gid1c-1*, and *gid1b-1 gid1c-1* mutants grown as in **d**. All data are means (±SEM) of *n* = 2-4 independent biological replicates. For **b** and **d-h**, DMSO (0.05 % v/v) treatment served as a control. Statistical analysis was performed using two-tailed Student’s *t*-test or one-way ANOVA, and letters denote significant differences with a Tukey’s *post hoc* test at *P*<0.05.

To distinguish between effects on GA biosynthesis versus signaling, we evaluated GA sensitivity in PAC-pretreated seedlings. Interestingly, while *cdc48a-4* and *npl4ab* responded similarly to WT at low GA concentrations, they exhibited a hypersensitive elongation response at higher doses (Fig. 4c), suggesting that the CDC48 complex modulates both GA metabolism and signaling. In agreement with a role in GA biosynthesis, *cdc48a-4* and CB-5083-treated seedlings showed increased levels of active GA, coupled to a depletion of biosynthetic intermediates (Fig. 4d; Supplementary Fig. S6c). Moreover, these seedlings also exhibited an accumulation of GA catabolites (Fig. 4d). Accordingly, the transcript levels of GA biosynthetic (*GA20OX1* and *GA3OX1*) and catalytic (*GA2OX1*) genes were induced in *cdc48a-4* or CB-5083-treated seedlings compared to WT (Fig. 4e). Elevated GAs are known to promote DELLA degradation. Consistently, the inhibition of CDC48A slightly reduced DELLA protein RGA levels (Fig. 4f; Supplementary Fig. S6d). However, the quintuple *della* mutant (*global della*) remained fully responsive to CB-5083-induced hypocotyl elongation (Fig. 4g; Supplementary Fig. S6e), suggesting that CDC48 regulatory impact on growth is largely independent of DELLA-mediated repression. Supporting this, seedlings overexpressing non-degradable DELLA variants also displayed WT-like elongation response when treated with CB-5083 (Fig. 4g).

To test whether the DELLA-independent function of CDC48 relies on GID1-mediated GA perception, we evaluated the hypocotyl response to CB-5083 in single and double mutants of the GA receptors. Interestingly, these seedlings displayed a reduced response to CDC48A pharmacological inhibition compared to WT controls, with *gid1a-1 gid1c-1* showing complete insensitivity to the treatment (Fig. 4h). Together, these results point to CDC48A acting through a GID1-mediated non-canonical GA signaling pathway.

### The CDC48-NPL4-UFD1 complex interacts with the GA receptor GID1

To elucidate the molecular mechanism through which CDC48A modulates GA signaling, we performed yeast two-hybrid (Y2H) assays to evaluate potential physical interactions between CDC48A-UN and the primary components of the GA perception and signaling. No binding was detected with RGA, whereas UFD1B showed binding to GID1A in yeast (Fig. 5a). In fission yeast, several point mutations in *UFD1* affecting conserved residues generate thermosensitive alleles (Nie et al. 2012) (Supplementary Fig. S8). Notably, PISA analysis of the three-dimensional structure of UFD1B-GID1A complex modelled using Alphafold3 identified a conserved glutamic acid (E) at the interaction interface corresponding to the site of these yeast mutations (Fig. 5b; Supplementary Table S1). To evaluate their functional relevance, we mutated E residues at positions 78 and 79 to alanine (A) in UFD1B, and observed that these substitutions markedly impaired the interaction with GID1A (Fig. 5c). Finally, we confirmed the association between CDC48A-UN and GID1 *in planta* using bimolecular fluorescence complementation (BiFC) in tobacco leaves. Indeed, co-expression of GID1A and NPL4B fused to complementary fragments of mCitrine resulted in a robust nuclear fluorescence signal (Fig. 5d). The binding of GID1A to CDC48A-UN was further validated via streptavidin-dependent gel shifts. In extracts from leaves expressing turboID-tagged versions of CDC48A or UFD1B, a fraction of GID1A exhibited a molecular weight shift upon streptavidin incubation (Fig. 5e). Collectively, our findings demonstrate that CDC48A-UN physically associates with the GA receptor GID1, highlighting a conserved motif within UFD1 as the primary interface required for this molecular recognition.

**Fig. 5.**
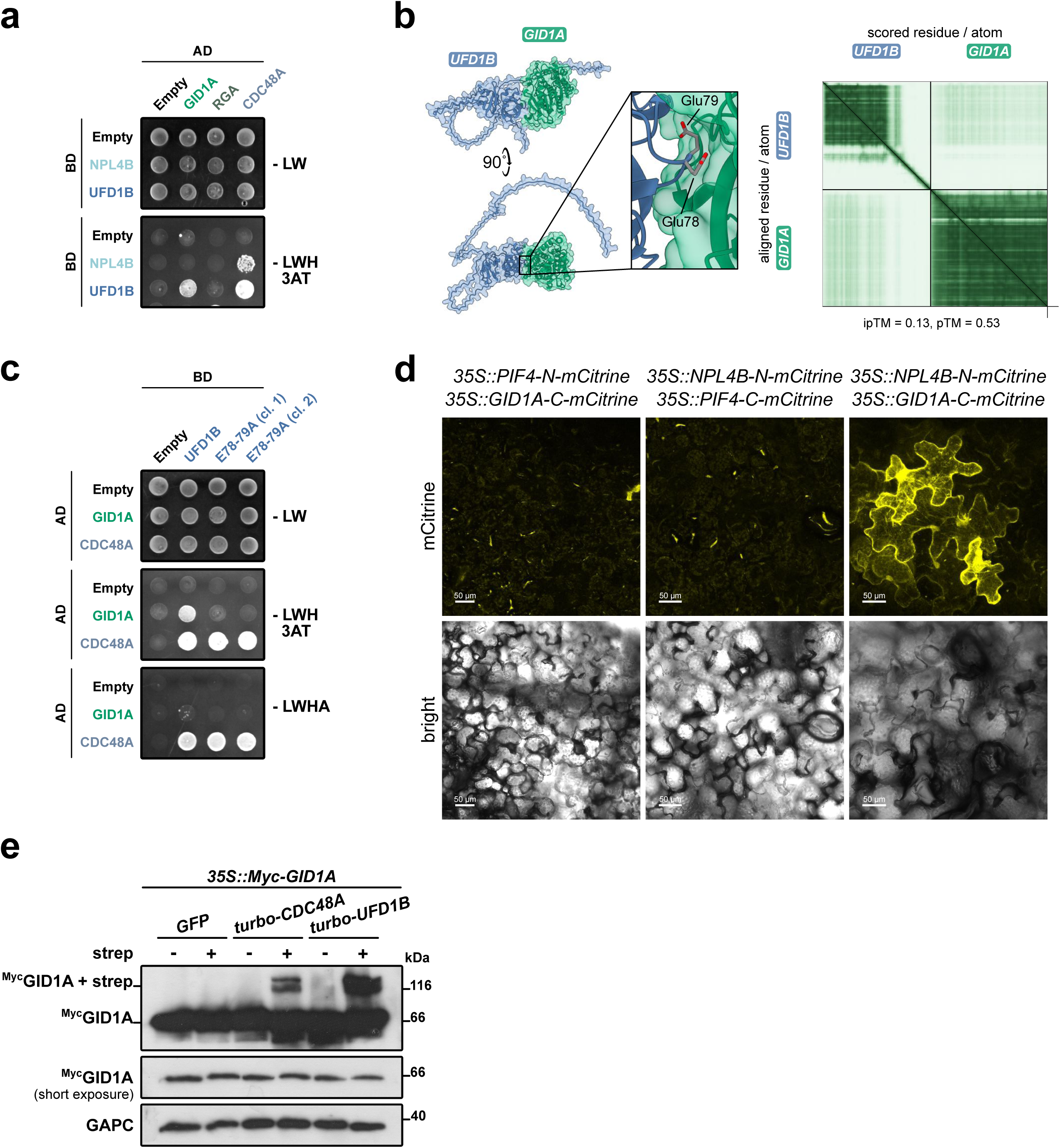
The co-factor UFD1 physically interacts with the GA receptor GID1. **a**, Y2H analysis of NPL4B or UFD1B with GID1A, RGA, or CDC48A. **b**, Left: AlphaFold-predicted structures of the interaction between UFD1B (blue) and GID1 (green), shown from two different orientations. The predicted template modeling (pTM) and interface predicted template modeling (ipTM) scores are indicated for each model. Right: Predicted aligned error (PAE) plot illustrating regions of high confidence (dark green) and low confidence (pale green) in the predictions. **c**, Y2H analysis of GID1A or CDC48A with UFD1B WT or the E78-79A mutant. **d**, Bimolecular fluorescence complementation (BiFC) analysis of the interaction between GID1A and UFD1B. Constructs expressing the indicated proteins fused to the N- or C-terminal of mCitrine were co-transformed into *Nicotiana benthamiana* leaf epidermal cells. Constructs expressing *PIF4* were used as negative controls. Scale bars, 50 µm. **e**, Immunoblot analysis of the streptavidin-induced electrophoretic mobility shift of ^Myc^GID1A. Protein extracts were obtained from tobacco leaves transiently co-expressing *35S::10xMyc-GID1A* with either *35S::NLS-GFP* (*GFP*), *35S::turboID-GFP-CDC48A* (*turbo-CDC48A*), or *35S::turboID-GFP-UFD1B* (*turbo-UFD1B*). GAPC served as loading control. For **a** and **c**, spotted yeast cells were grown for 3 d on control medium (-LW; lacking leucine and tryptophan) or selective medium (-LWH + 3AT: lacking leucine, tryptophan, and histidine and supplemented with 1 mM 3-aminotriazole; -LWHA: lacking leucine, tryptophan, histidine, and adenine). Constructs fused to the Gal4 activation domain (AD) or Gal4 DNA-binding domain (BD) are indicated. Interaction with CDC48A served as positive control.

### The CDC48-UFD1-NPL4 complex promotes GID1 degradation

Since CDC48-UN is best known for targeting proteins for degradation (Meyer and van den Boom 2023), we next evaluated whether it might regulate GID1 stability. Transient overexpression of *^Myc^GID1A* in tobacco leaves revealed that co-expression of *CDC48A* was sufficient to decrease ^Myc^GID1A protein levels compared to WT controls (Fig. 6a). Notably, overexpression of the *cdc48a-4* mutant version restored GID1A protein abundance to WT levels (Fig. 6a). These results suggest that CDC48A modulates GID1A protein abundance, likely by promoting its degradation. Furthermore, live-cell imaging demonstrated that *CDC48A* overexpression significantly depleted the nuclear pool of ^mCherry^GID1A (Fig. 6b,c). To further confirm the role of CDC48A in GID1 turnover, we grew reporter lines expressing GUS-tagged GA receptors controlled by their endogenous promoters under blue light in the presence or absence of CDC48A inhibitor. Although *GID1C* transcripts were reduced in *cdc48a-4* mutants and in CB-5083-treated seedlings, GUS activity in the hypocotyl was increased upon CDC48A inhibition (Fig. 6d,e). In contrast to GID1C, seedlings expressing GUS-tagged GID1B showed no detectable histochemical staining in the hypocotyl under blue light conditions (Supplementary Fig. S9). These results suggest that CDC48A-UN is required for efficient proteasomal processing of the GA receptor GID1A and GID1C. Altogether, our results reveal an uncharacterized role of CDC48A, together with NPL4 and UFD1, in integrating light and hormonal cues through protein homeostasis to regulate photomorphogenic development.

**Fig. 6.**
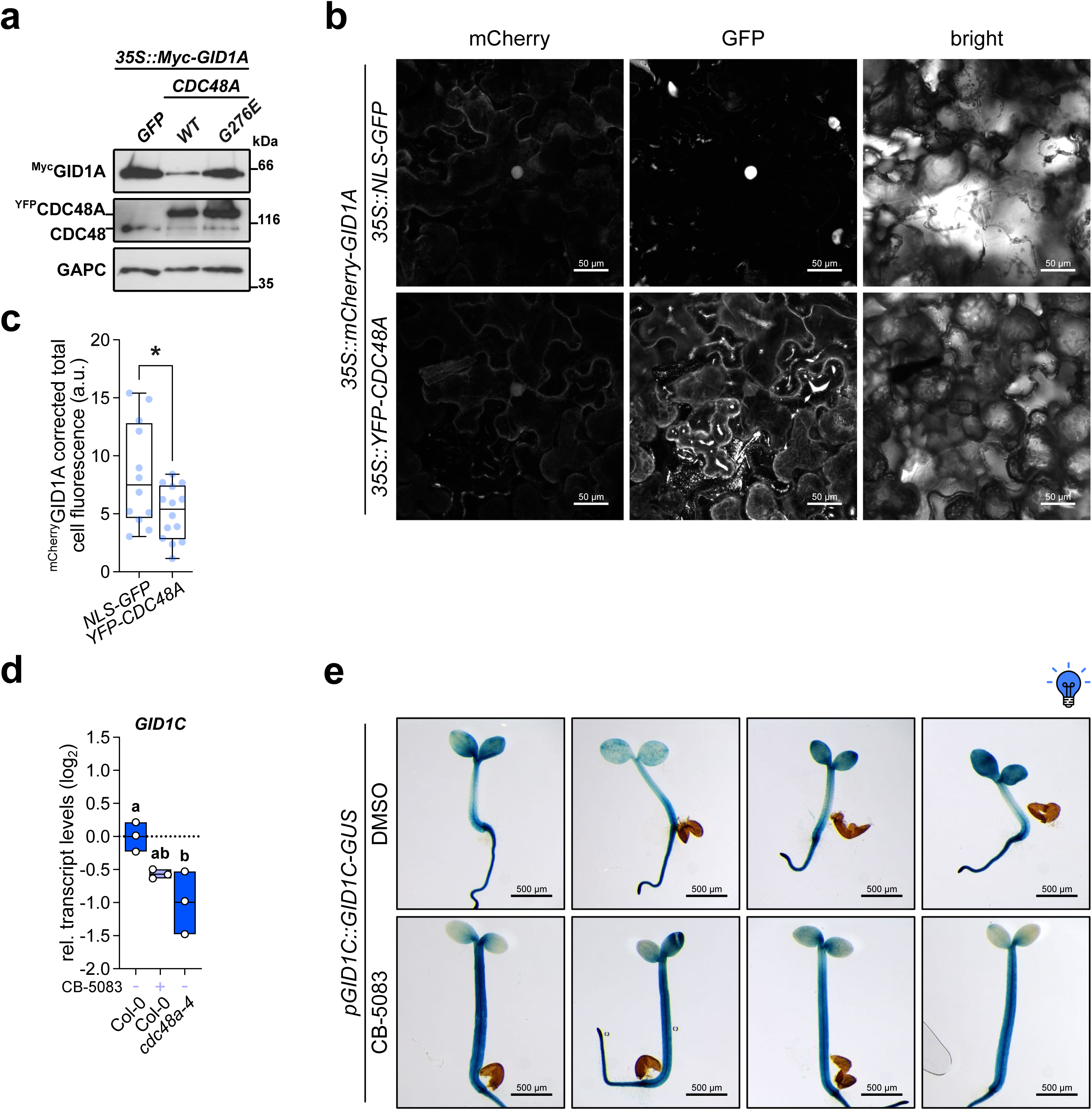
The CDC48A-NPL4-UFD1 complex promotes GID1 turnover. **a**, Immunoblot of ^Myc^GID1A and CDC48 from tobacco leaves co-transformed with *35S::10xMyc-GID1A* and *35S::NLS-GFP* (*GFP*), *35S::YFP-CDC48A* (*CDC48A WT*), or *35S::YFP-CDC48A G274E* (*CDC48A G274E*). GAPC served as loading control. **b-c**, Representative sum-intensity Z-projections (**b**) and quantification (**c**) of fluorescence signal from tobacco leaves co-transformed with *35S::mCherry-GID1A* and *35S::NLS-GFP* or *35S::YFP-CDC48A*. Sum-intensity Z-projections are shown. Scale bars, 50 µm. **d**, *GID1C* transcript levels quantified by RT-qPCR in Col-0 wild-type and *cdc48a-4* seedlings grown under continuous blue light for 4 d at 22 °C, in the presence or absence of the CDC48 activity inhibitor CB-5083 (1 µM). DMSO (0.05 % v/v) treatment served as a control. Data were normalized to *ACTIN2/8* transcript levels and are shown on a log_2_ scale relative to Col-0 grown in DMSO, which was set to zero. Data are presented as floating bar plots, with the line indicating the mean of *n* = 3 independent biological replicates and individual values shown. Different letters indicate significant differences among means as determined using one-way ANOVA followed by Tukey’s *post-hoc* test (*P*<0.05). **e**, Representative images of GUS staining for transformed Col-0 seedlings expressing *GID1C* fused to the *GUS* reporter under the control of *GID1C* endogenous promoter (*pGID1C::GID1C-GUS*), grown as in **d**. Scale bars, 500 µm. For **a** to **c**, tobaccos were grown under long-day conditions at 22 °C.

## Discussion

Photomorphogenesis is a light-driven developmental program that orchestrates major morphological and physiological transitions in plants. Because light modulates the stability of regulatory and effector proteins, precise control of protein homeostasis is essential for appropriate responses (Zhou and Deng 2025). Here, we identified the segregase CDC48 as a novel modulator of blue light-mediated photomorphogenesis that acts through gibberellin signaling.

Similar to other eukaryotes, CDC48A localizes to the cytoplasm, the plasma membrane, and the nucleus in plants, with specific stimuli (such as cell cycle and virus infection) triggering CDC48A subcellular recruitment (Park et al. 2008; Niehl et al. 2012). While the nuclear pool of CDC48 is depleted during the immune response (Inès et al. 2025), our data revealed that blue light acts as a novel environmental cue that triggers CDC48 multicompartmental redistribution (Fig. 1; Supplementary Fig. S1). Although CRY1 promotes the nuclear import of Heat shock factor A1d (HsfA1d) (Gao et al. 2023), CDC48A maintained its subcellular distribution in *cry1-304* (Supplementary Fig. S1b), suggesting that blue light-dependent CDC48A accumulation involves compensatory mechanisms or additional photoreceptors, such as CRY2. The molecular drivers of these dynamics and the potential of the light-responsive accumulation across species warrant further investigation.

During photomorphogenesis, blue light perception restricts hypocotyl growth while concurrently stimulating cotyledon expansion and chloroplast biogenesis (Liscum and Hangarter 1991). We found that CDC48 segregase activity is required to repress hypocotyl elongation under blue light (Fig. 2; Supplementary Fig. S2). Notably, the synergistic effect of the different *cdc48* mutants suggests that all three isoforms contribute to photomorphogenesis, albeit to different extents (Fig. 2h; Supplementary Fig. S2d). Note that the marked reduction in hypocotyl growth in these lines following CB-5083 application likely results from the inhibition of essential CDC48 functions, which might impact seedling viability at higher concentrations or in sensitized backgrounds (Fig. 2g,h). Furthermore, our genetic analysis indicated that CDC48 functions through the canonical photomorphogenic pathway downstream of blue light perception, requiring functional COP1 and HY5 (Fig. 2; Extended Data Figs. 2 and 3). Additionally, the elongated phenotype of CDC48A-deficient lines was strictly PIF-dependent (Fig. 2j). Although PIFs typically regulate hypocotyl growth through auxin(Krahmer and Fankhauser 2024), we demonstrated that CDC48A functions independently of auxin metabolism, transport, and signaling (Supplementary Fig. S5). This aligns with models where PIF4/5 promote elongation under blue light by regulating cell-wall-modifying proteins rather than auxin levels or sensitivity (Pedmale et al. 2016).

CDC48 recruits a specialized set of cofactors that confer substrate specificity, such as the NPL4-UFD1 adapter module for processing of ubiquitin-modified targets (Meyer and van den Boom 2023). Unlike the obligate heteromeric complexes found in animals and yeasts, plant NPL4 and UFD1 associate with CDC48 independently, yet their combined interaction enhances substrate unfolding efficiency(Huntington et al. 2026). Consistent with this cooperative mechanism, we found that both cofactors are essential for the CDC48A-mediated photomorphogenic response (Fig. 3). Since these adapters also coordinate heterochromatin decondensation and plastid proteostasis under stress (Mérai et al. 2014; Li et al. 2022), our data reinforce the model that plant NPL4 and UFD1 function cooperatively as in other eukaryotes. However, whether these cofactors possess distinct, independent roles in plants remains to be determined.

Similar to auxin, gibberellins stimulate hypocotyl elongation under different environmental cues. We demonstrated that CDC48A-mediated growth regulation requires a functional GA pathway, with CDC48A activity modulating GA biosynthesis and signaling (Fig. 4; Supplementary Fig. S6). While inactivation of CDC48A induced the expression of GA biosynthetic genes and increased bioactive GA pools, catabolic genes and catabolite levels were also upregulated (Fig. 4d,e). These results suggest a robust metabolic feedback to counteract hormone overproduction or excessive signaling, which aligns with responses observed in cytokinin-deficient plants that coordinate both GA biosynthesis and degradation to ensure metabolic stability (Werner et al. 2025).

Besides the canonical GA-GID1-DELLA module, several studies have shown that GA modulates plant development or environmental responses through DELLA-independent pathways. For instance, the transcription factor SPATULA (SPT) promotes fruit growth through GID1-mediated GA perception while bypassing DELLA repression (Fuentes et al. 2012). Given that SPT does not physically associate with GID1, it likely requires an unidentified molecular bridge or cofactor. GA stimulates GID1-mediated turnover of NITROGEN-MEDIATED TILLER GROWTH RESPONSE 5, which facilitates the nitrogen-dependent silencing of shoot branching inhibitors by the Polycomb Repressive Complex 2 (Wu et al. 2020). Rapid signaling events, such as GA-induced increases in cytosolic Ca^2+^, also operate independently of the DELLA pathway (Okada et al. 2017). Similarly, we found that the CDC48 segregase represses hypocotyl growth through a DELLA-independent GID1 signaling pathway (Fig. 4), potentially by modulating the GID1-mediated degradation of an unidentified negative GA regulator. Since CDC48 is implicated in endoplasmic reticulum homeostasis (Müller et al. 2005; Zang et al. 2020; Schoberer et al. 2024), a major intracellular Ca^2+^ reservoir, it is tempting to speculate that this GA-mediated growth regulation converges on Ca^2+^-dependent signaling. However, the molecular mechanisms of the DELLA-independent pathway/s require further investigation.

CDC48 is a core protein quality control segregase that facilitates proteasomal degradation, using the UFD1-NPL4 adapter module for processing of ubiquitin-modified targets. In plants, CDC48-UN likely promotes the extraction and subsequent turnover of the conserved inner nuclear membrane protein SUN1 (Sad1 and UNC84 Domain Containing 1), and regulates the ubiquitin-dependent degradation of intrachloroplast proteins, such as RbcL (ribulose-bisphosphate carboxylase large subunit) and AtpB (ATP synthase subunit beta) (Huang et al. 2020; Li et al. 2022; Calvanese et al. 2025). We expanded this regulatory repertoire in plants by demonstrating that CDC48A-UN interacts with the GID1 receptors to modulate GA signaling under blue light (Fig. 5). Recent evidence in yeast indicates that mutating a conserved leucine within the UT3 domain of Ufd1 selectively disrupts ubiquitin binding without compromising Cdc48-Npl4 complex assembly. By mutating conserved residues in UFD1B at the GID1A interaction interface, we abolished their association without disrupting CDC48A binding (Fig. 5b,c). This confirms that GA receptor recruitment can be structurally uncoupled from core segregase assembly. Moreover, it suggests that plants may have co-opted a pre-existing structural module within UFD1 to facilitate a novel hormone-signaling function. Whether this interface is uniquely dedicated to GA perception or serves as a broader recognition hub for additional substrates warrants deeper exploration.

While GID1-mediated DELLA degradation upon GA perception is well-characterized(Shani et al. 2024), the mechanisms governing receptor stability remain poorly understood. Beyond the GA receptor RING E3 ubiquitin ligase (GARU), which mediates ubiquitin-dependent GID1 turnover (Nemoto et al. 2017), our results expanded this regulatory landscape by demonstrating that CDC48 also promotes GID1 degradation (Fig. 6). This raises the question of whether CDC48 operates downstream of GARU or through an alternative E3 ligase. Interestingly, the CDC48-interacting factor PUX1 associates with and stabilizes GID1, although the precise molecular mechanism remains elusive (Hauvermale et al. 2022). One compelling hypothesis is that PUX1 promotes GID1 accumulation by inhibiting CDC48 activity. However, in contrast to our results, DELLAs are significantly degraded in *pux1* mutants (Hauvermale et al. 2022). Furthermore, *pux1* maintained WT hypocotyl elongation under blue light (Supplementary Fig. S7). Thus, the PUX1-mediated regulation of GID1 may be either context-specific or redundant with other PUX members during photomorphogenesis.

Despite CDC48A inactivation leading to higher GA levels and GID1 stabilization, DELLA proteins exhibited only a slight reduction under blue light (Fig. 4f). While this could reflect non-proteolytic DELLA downregulation, as previously reported (Ariizumi et al. 2008; Fukazawa et al. 2015), CDC48-mediated growth repression remained largely independent of DELLAs (Fig. 4g). This divergence from the canonical model might be attributed to a deficient GID1-DELLA association. Supporting this, activated CRY1 antagonizes DELLA recruitment by GID1, effectively uncoupling receptor abundance from downstream protein degradation (Xu et al. 2021; Yan et al. 2021; Zhong et al. 2021).

In conclusion, we propose a model in which blue light-mediated subcellular partitioning of CDC48-UN is critical to modulate GA homeostasis and trigger GID1 receptor turnover. This regulatory axis activates a non-canonical GA signaling pathway through an unidentified effector, which facilitates photomorphogenesis by establishing a cellular environment that antagonizes PIF-mediated hypocotyl elongation (Fig. 7). This mechanism reveals a novel layer of hormonal regulation, in which the CDC48-UN segregase functions as an upstream gatekeeper of GA signaling output during blue light-induced photomorphogenesis.

**Fig. 7.**
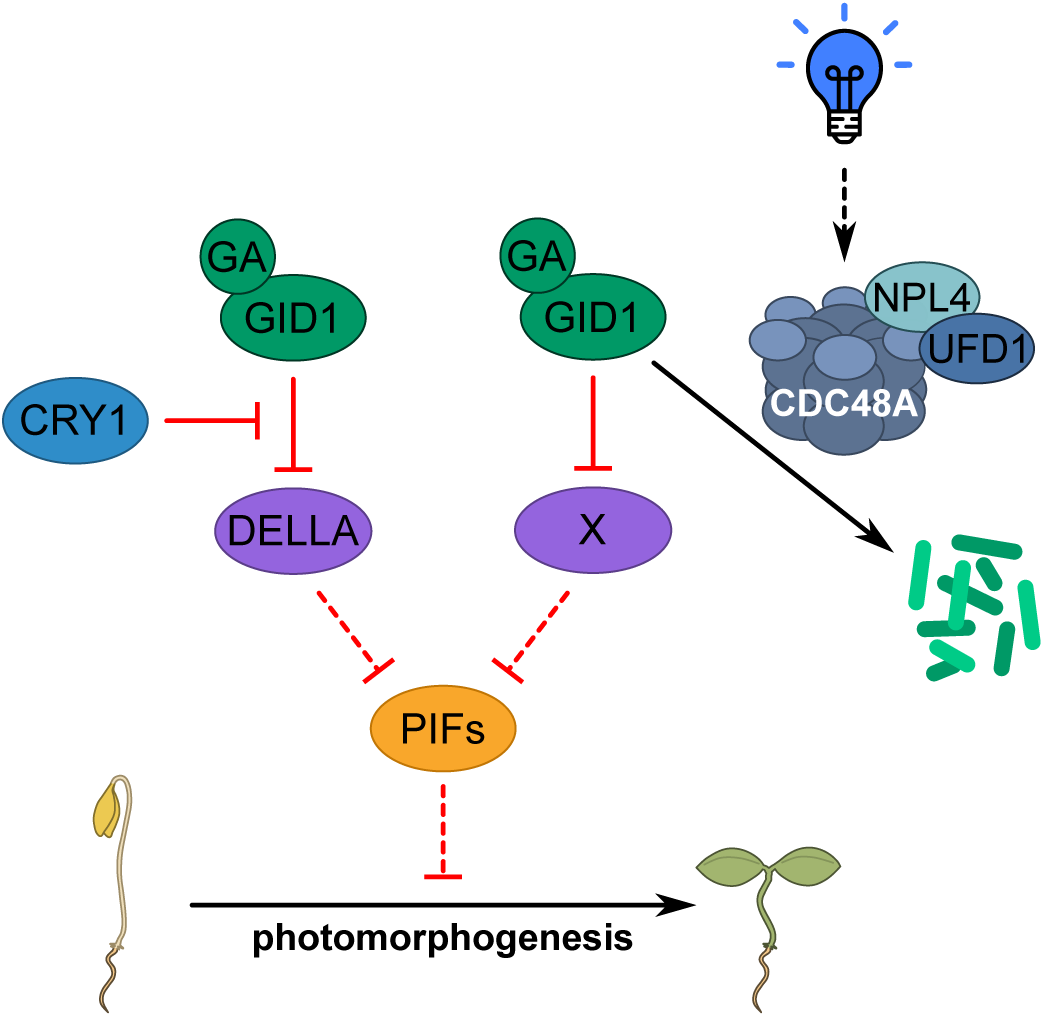
Proposed model for the role of the CDC48A-UFD1-NPL4 complex in blue light-mediated photomorphogenesis. Beyond the traditional GA signaling pathway, blue light induces a specific subcellular shift in CDC48-UN localization, which facilitates the recognition and subsequent turnover of the GA receptor GID1. Operating independently of the canonical GA-GID1-DELLA signaling module, this CDC48-mediated reduction in GID1 abundance results in the stabilization of an unidentified negative regulator, which actively restricts hypocotyl elongation.

## Materials and Methods

### Plant material

Wild-type and other lines of plants used in this study were of *Arabidopsis thaliana* L. Heynh. ecotype Columbia (Col-0) or Landsberg *erecta* (L*er*). Seeds of the following genotypes were obtained from the Arabidopsis Biological Resource Center (ABRC; https://abrc.osu.edu): *axr1-3* (CS57504), *axr1-12* (CS3076), *ga3ox1-3 ga3ox2-1* (CS6944), *pif1-2* (SALK_071677), *pif3-1* (CS66042), *pif4-2* (CS66043), *pif5-3* (CS66044), *pif1-1 pif3-7 pif4-2 pif5-3* (*pifq*; CS66049), *pux1-3* (CS923389), and *yucca6-1D* (CS67234) in the Col-0 ecotype background, and the plants overexpressing either *RGA* (*35S::TAP-RGA*Δ*17*; CS16292) or *GAI* (*35S::TAP-GAI*Δ*17*; CS16294) in the L*er* ecotype. The *cdc48a-4 cdc48b-3*, *cdc48a-4 cdc48b-2*, *cdc48a-4 cdc48c-1*, *cdc48b-3 cdc48c-2*, and *cdc48a-4 cry1-304* double mutants were generated by crossing the corresponding single mutants and subsequently confirmed by genotyping (primer sequences listed in Supplementary Table S2). The *cdc48b-3* (GABI_104F08), *cdc48c-2* (SALK_102955), *cdc48c-3* (SALK_123409), *gid1a-1* (SALK_142767), *gid1b-1* (SM_3_30227), *gid1c-1* (SALK_023529), *gid1a-1 gid1b-1*, *gid1a-1 gid1c-1*, *gid1b-1 gid1c-1*, *pGID1B::GID1B-GUS*, *pGID1C::GID1C-GUS*, and *DR5::GUS* in Col-0 ecotype, and the quintuple *della* mutant (*gai-t6 rga-t2 rgl1-1 rgl3-1 rgl2-1*) and *pRGA::GFP-RGA* seedlings in L*er* ecotype were previously described (Ulmasov et al. 1997; Silverstone et al. 2001; Suzuki et al. 2009; Gallego-Giraldo et al. 2014; Blanchard et al. 2025; Garro et al. 2025). The mutant seeds *cdc48a-4* (CS69995), *cdc48b-2* (SAIL_566_E10), *cdc48c-1* (SAIL_1182_E09), *cdc48D-1* (SALK_074372), and *cdc48b-2 cdc48c-1*(Copeland et al. 2016) were generously provided by Prof. Xin Li from the University of British Columbia, Vancouver, Canada. The *npl4a npl4b* (*npl4ab*), *ufd1b ufd1c* (*ufd1bc*), *proCDC48A::EYFP-CDC48A #3* (*pCDC48A::YFP-CDC48A*), and *35S::NPL4B-GFP #7* (*35S::NPL4B-GFP*) (Li et al. 2022) seeds were kindly supplied by Prof. Rongcheng Lin from the Institute of Botany, Chinese Academy of Sciences, Beijing, China. The seeds bearing estradiol-inducible *CDC48-WT* and *CDC48-DN* constructs (Ling et al. 2019) were generously provided by Prof. Paul Jarvis from the University of Oxford, Oxford, United Kingdom. The mutant seeds *pif7-1* (CS68809) (Leivar et al. 2008a), *pif4-2 pif5-3* (*pif45*) (Lorrain et al. 2008), *pif4-2 pif5-3 pif7-1* (*pif457*) (Willige et al. 2021), and *cry1-17* (Ahmad et al. 1995) were kindly provided by Dr. Yogev Burko from the Agricultural Research Organization, Bet Dagan, Israel. The *cry2-1 pUBQ10::9xMyc-6xHis-3xFlag-CRY2* (*CRY2*-ox) (Pedmale et al. 2016) seeds were generously provided by Dr. Ullas V. Pedmale from the Cold Spring Harbor Laboratory, New York, USA. The *cop1-4*, *cop1-6*, *cry1-304*, *pif4-101*, *pif3-3 pif4-2*, *pif4-101 pif5-3*, *pif3-3 pif4-2 pif5-3*, and *pif4-101 pif5-3 pif7-1* seeds were generously supplied by Dr. Jorge J. Casal from Instituto de Investigaciones Fisiológicas y Ecológicas Vinculadas a la Agricultura (IFEVA), Buenos Aires, Argentina.

### Plant growth conditions and treatments

Arabidopsis plants were grown on soil in a 21-23 °C growth chamber under a long-day photoperiod (16 h light, 110 μmol m^-2^ s^-1^/8 h dark). Seeds for hypocotyl length evaluation were surface-sterilized by treatments with 70% EtOH and 10% sodium hypochlorite, washed, and stratified at 4 °C for 3 days to obtain homogeneous germination. Unless stated otherwise, seedlings were grown in Petri dishes containing half-strength Murashige and Skoog (½MS) basal medium (Murashige and Skoog 1962) supplemented with vitamins (PhytoTechnology Laboratories) and 0.9% agar, at 21-23 °C under long-day conditions (16 h light, 110 μmol m^-2^ s^-1^/8 h dark) for 4 days. For light treatments, continuous monochromatic blue and red lights were generated from blue LEDs (λ max = 469 nm) and red LEDs (λ max = 680 nm), respectively. For treatments, seeds were grown in ½MS medium containing 0.1 to 5 µM CB-5083 (Cayman Chemicals, Ann Harbor, MI, USA), 0.5 µM picloram (PIC; Duchefa Biochemie, Haarlem, the Netherlands), 0.5 µM N-1-naphthylphthalamic acid (NPA;), 0.1 to 10 µM GA_3_ (Duchefa Biochemie, Haarlem, the Netherlands), or 1 mM paclobutrazol (PAC; Sigma-Aldrich, St. Louis, MO, USA) at 21-23 °C under the different light conditions for 4 days. The CB-5083, GA_3_, and PAC stock solutions were prepared according to the manufacturer’s recommendation. Relative response was defined as the ratio of hypocotyl length in CB-5083-treated seedlings to that of the same genotype exposed to DMSO.

### Genetic constructs

Cloning was performed using standard molecular biology procedures or the Gateway™ recombination cloning system (Invitrogen™). For BiFC assays, the full-length coding region of *GID1A*, and *NPL4B* cloned into *pENTR3C* was recombined into *pAS-054* and *pAS-059* (Hellens et al. 2000) to generate C-terminal fusions to the N- and C-terminal moieties of mCitrine. The *pAS-054* and *pAS-059* constructs bearing the *PIF4* coding sequence were previously described (Ferrero et al. 2019). For yeast two-hybrid (Y2H) assays, coding sequences were amplified using gene-specific oligonucleotides (Supplementary Table S3) and cloned into the *pGADT7* or *pGBKT7* vectors (Clontech™). To generate overexpression constructs, the *CDC48A*, *UFD1B,* and *CRY1* coding sequences were amplified from Arabidopsis RNA and cloned into the *pENTR3C* using the restriction sites indicated in Supplementary Table S3. The inserts were subsequently recombined using Gateway™ technology into the destination vectors *pB7WGY2* (Karimi et al. 2005), *pGW-GFP-TurboID-GOI* (Addgene #209381) (Tan et al. 2024), or *pFAST-R06* (Shimada et al. 2010), generating the constructs *35S::YFP-CDC48A*, *35S::turboID-GFP-CDC48A* or *35S::turboID-GFP-UFD1B*, and *35S::GFP-CRY1* (*CRY1*-ox), respectively. Similarly, the *GID1A* coding sequence was amplified, cloned into *pENTR3C*, and subsequently recombined into *pJV-117* (Hellens et al. 2000) or *pGWB421* (Addgene #74815) (Nakagawa et al. 2007) to produce the vectors *35S::mCherry-GID1A* and *35S::10xMyc-GID1A*, respectively. The dominant-negative *35S::turboID-GFP-CDC48A E581Q* (*CDC48A EQ*) variant was created by introducing the E581 to Q mutation into the entry clone before the recombination with *pGW-GFP-TurboID-GOI*. Plasmids carrying point mutations were generated by PCR-based site-directed mutagenesis using specific primers (Supplementary Table S3). A complete list of plasmids used in this study is provided in Supplementary Table S4. Plasmid maps and DNA sequences are available upon request.

### Plant transformation

Arabidopsis plants were stably transformed via floral dip using *Agrobacterium tumefaciens* LBA4404 harboring the specific constructs (Clough and Bent 1998). Transgenic plants were selected on ½MS medium supplemented with the appropriate selective agent (ammonium glufosinate 1 ml l^-1^, or kanamycin 25 mg l^-1^). Seeds were surface-sterilized, stratified for 2 days at 4 °C, and then germinated in a growth chamber at 21-23 °C. When ammonium glufosinate was used, selection was performed directly in soil. For each construct, three to four independent positive lines were propagated, and homozygous T3 or T4 plants carrying a single insertion, as determined by Mendelian segregation, were used for further analyses. For transient localization assays, *Nicotiana benthamiana* leaves were infiltrated with *Agrobacterium tumefaciens* LBA4404 carrying the indicated constructs, together with *A. tumefaciens* harboring the silencing suppressor p19 (de Felippes and Weigel 2010). Infiltrated plants were incubated under the indicated light conditions, and samples were collected 48 h post-infiltration for analysis by confocal laser scanning microscopy (Leica TCS SP8).

### Measurement of hypocotyl and cell length

As previously reported (Capella et al. 2015), hypocotyl length was measured as the distance from the most basal root hair to the “V” formed by the cotyledons. Each biological replicate corresponds to the mean hypocotyl length of at least 15 seedlings grown on the same plate. Graphs represent mean values obtained from 3-4 plates per treatment and/or genotype. Cell length was assessed using confocal microscopy, with each biological replicate representing the average length of 8-10 cells from an individual seedling. Hypocotyl and cell length were determined using Fiji/ImageJ software (Schindelin et al. 2012).

### Histochemical GUS staining

*In situ* GUS staining was performed as previously described (Jefferson et al. 1987). Seedlings were grown on ½MS basal medium with or without 1 µM CB-5083 at 21-23 °C under continuous blue light conditions for 4 days. Seedlings were immersed in GUS staining buffer (1 mM 5-bromo-4-chloro-3-indolyl-β-glucuronic acid solution in 100 mM sodium phosphate pH 7, 0.1% (v/v) Triton X-100, 2 mM potassium ferrocyanide, and 2 mM potassium ferricyanide), subjected to vacuum infiltration for 5 min, and incubated at 37 °C overnight. Chlorophyll was removed from stained tissues by immersing the seedlings in 70% ethanol.

### Quantification of gibberellins (GAs)

GAs were quantified as previously described (Seo et al. 2011). Briefly, metabolites from seedlings grown under continuous blue light for 4 d at 22 °C were extracted using 80% methanol containing 1% acetic acid. The extracts were purified by consecutive passage through HLB (reverse phase), MCX (cationic exchange), and WAX (anionic exchange) SPE columns (Oasis, 30 mg; Waters). GAs were separated via reverse-phase chromatography (2.6 µm Accucore RP-MS column; ThermoFisher Scientific) using an acetonitrile gradient (5% to 50%) with 0.05% acetic acid at a flow rate of 400 µL/min. Mass spectrometry analysis was performed using electrospray ionization (ESI) and targeted-SIM on a Q-Exactive Orbitrap spectrometer (ThermoFisher Scientific). Deuterated [17,17-2H] GA_1_, GA_4_, GA_8_, GA_9_, GA_12_, GA_15_, GA_19_, GA_24_, GA_34_, GA_51_, and GA_53_ were used as internal standards for quantification.

### Yeast two-hybrid assays

Yeast manipulations were performed following standard transformation and mating protocols. Cells were grown at 30 °C in YPD medium (1% yeast extract, 2% peptone, and 2% glucose) or synthetic complete medium lacking individual amino acids to maintain selection for transformed plasmids. The *Saccharomyces cerevisiae* AH109 and Y187 (Clontech^TM^) strains were transformed with the indicated *pGBKT7* or *pGADT7* constructs, respectively, followed by mating. Spotting assays were performed on control medium (-LW) or selective medium with increased stringency (-LWH, -LWH + 1 mM 3-aminotriazol (3AT), -LWHA). Plates were incubated for 3 days at 30 °C unless indicated otherwise.

### Fluorescence microscopy

For localization analyses, *pCDC48A::YFP-CDC48A* and *35S::NPL4B-GFP* seedlings were grown on ½MS basal medium at 21-23 °C under the different light conditions for 4 days. In addition, transient expression assays were carried out in *Nicotiana benthamiana* leaves using *35S::GFP-CDC48A* or *35S::GFP-UFD1B* constructs to confirm subcellular localization. For quantification of nuclear fluorescence intensity, *pRGA::GFP-RGA* seedlings were grown on ½MS basal medium in the presence or absence of 1 µM CB-5083 at 21-23 °C under continuous blue light conditions for 4 days. For analysis of GID1A protein abundance, the construct *35S::mCherry-GID1A* was transiently co-expressed with either *35S::YFP-CDC48A* or *35S::NLS-GFP* in *N. benthamiana* leaves. At least 10 nuclei were analyzed for each transformation. A minimum of 100 nuclei from at least 8 independent biological samples were measured for each treatment. Corrected total cell fluorescence intensity was calculated as follows: CTCF = [integrated density – (area of selected cell × mean fluorescence of background fluorescence)]/1000.

Images, overlays, and Z-stacks were acquired with a confocal inverted microscope (Confocal LEICA TCS SP8), using a 10× objective. For GFP-tagged proteins, excitation was performed using a 488 nm laser line (50% intensity), and emission was collected between 493-522 nm using bandpass filters. For YFP-tagged proteins, excitation was performed using a 514 nm laser line (30% intensity), and emission was collected between 518-567 nm using bandpass filters. For mCherry-tagged proteins, excitation was performed using a 552 nm laser line (50% intensity), and emission was collected between 580-650 nm using bandpass filters. All images were captured using identical microscope settings. Subsequent processing and analyses of the images were performed in Fiji/ImageJ (Schindelin et al. 2012).

### Bimolecular fluorescence complementation (BiFC) assays

The genetic constructs to express GID1A, NPL4B, and PIF4 fused to the N- and C-terminal fragments of mCitrine were introduced into *A. tumefaciens* LB4404 and co-infiltrated into *N. benthamiana* leaves in different combinations in a 1:1 ratio as previously described (de Felippes and Weigel 2010). Transformed plants were maintained under long-day conditions at 21-23 °C for 48 h before imaging using a confocal laser scanning microscope (TCS SP8; Leica Microsystems, Manheim, Germany). Fluorescence intensity of mCitrine was quantified with Fiji/ImageJ (Schindelin) using corrected total cell fluorescence (CTCF = {integrated density − [area of selected cell × mean fluorescence of background fluorescence]}/1000).

### RNA isolation and analysis

Transcript levels were analyzed by RT-qPCR using RNA extracted from seedlings grown on ½MS medium at 21-23 °C under continuous blue light for 4 days. For each genotype, at least 80 seedlings were pooled per sample, and 3-4 biological replicates were included per experiment. The total RNA was purified from seedlings as previously described (Chang et al. 1993). One μg of RNA was reverse-transcribed using oligo(dT)_18_ and M-MLV reverse transcriptase II (Promega). Quantitative real-time PCR (qPCR) assays were performed using a StepOne equipment (Applied Biosystems); each reaction contained a final volume of 20 μL that included 2 μL of SyBr green (4×), 8 pmol of each primer, 2 mM MgCl_2_, 10 μL of a 1/20 dilution of the RT reaction, and 0.1 μL of Taq Polymerase (Invitrogen), using standard protocols (40 to 45 cycles, 60 °C annealing). Fluorescence was quantified at 72 °C. Specific primers for each gene were designed and are listed in Supplementary Table S5. The expression levels were normalized using *ACTIN2/8* (AT3G18780/AT1G49240) as a reference gene, and quantification was carried out using the ΔΔCt method (Pfaffl 2001) relative to Col-0 seedlings grown at 22 °C.

### RNA-seq analysis

We reanalyzed public RNA-Seq data from 6-day-old Col-0 wild-type and *cry1 cry2* seedlings grown in continuous darkness, blue, or red light (Jiang et al. 2023). RNA-seq reads were analyzed on the Galaxy platform (Abueg et al. 2024). Briefly, adapter sequences were removed from the reads using Trimmomatic (Galaxy Version 0.39+galaxy2). The quality-filtered reads were aligned to the Arabidopsis reference genome (TAIR10) using RNA STAR (Galaxy Version 2.7.11b+galaxy0), employing a maximum intron length of 20,000 bp and guided by the gene and exon annotation from Araport V11. The read counts on each gene were then calculated using featureCounts (Galaxy Version 2.1.1+galaxy0). Differential expression analysis was performed using edgeR (Galaxy Version 3.36.0+galaxy5). Bigwig coverage files were generated using bamCoverage (Galaxy Version 3.5.4+galaxy0), and subsequently used for plotting with pyGenomeTracks (Galaxy Version 3.8+galaxy2). RNA-seq data were previously deposited in the Gene Expression Omnibus (GEO) database under the accession numbers GSE226927 and GSE246071.

### Alphafold protein structure prediction

The modeling of the UFD1B-GID1A complex was performed using Alphafold3 (version 3.0.0). First, the sequences were converted with Alphafold3_tools from fasta to a json file with the required structure (Park et al. 2025). Then, Alphafold3 was run with the default parameters on an NVIDIA A30 graphics card, and the top-ranked predicted model was used for further analysis. was generated using. Confidence summary statistics, such as ipTM and pTM, as well as the Predicted aligned error (PAE), were extracted from the Alphafold3 output. Protein structure visualization and the PAE plot were generated using UCSF ChimeraX (version 1.11) software and PAE viewer, respectively (Pettersen et al. 2021; Elfmann and Stülke 2023).

### Immunoblotting

Total protein extraction was performed as previously reported (Tonetti et al. 2025). Briefly, proteins from 20 mg FW of homogenized frozen tissue were extracted with 200 μl of 125 mM Tris–HCl pH 6.8, 2% (w/v) SDS, 20% (v/v) glycerol, 1.4 M 2-mercaptoethanol, and 0.05% (w/v) Bromophenol Blue. Samples were vortexed and incubated at 95 °C for 5 min with agitation, then cooled to room temperature and centrifuged at 20,000 g for 10 min to remove insoluble debris. Solubilized proteins were resolved on self-made 10% gels, transferred onto PVDF membranes (polyvinylidene fluoride membranes, GE Healthcare), and analyzed by standard immunoblotting techniques using specific antibodies. For streptavidin-induced gel shift assays, 10 µg of streptavidin was added to each sample and incubated for 5 min before loading into the gel. GAPC was used as a loading control.

### Antibodies

Polyclonal Cdc48 (1:5,000) antibody was raised in rabbits and has been described previously (Richly et al. 2005). Monoclonal antibodies directed against the c-Myc epitope (1:1,000; 9E10) and against GFP (1:1,000; B-2) were purchased from Sigma and Santa Cruz Biotechnology, respectively. Rabbit polyclonal Anti-GAPC1/2 (1:10,000; AS15 2894) was obtained from Agrisera. Secondary antibodies fused to HRP were used for detection (goat anti-mouse HRP 1:10,000, Thermofisher Scientific, 31430; goat anti-rabbit HRP 1:10,000, Thermofisher Scientific, 31460).

### Statistics and reproducibility

All experiments were performed at least twice with similar outcomes, and each figure panel displays representative results from these repetitions. Analyses of the variance were performed, and pairwise differences were evaluated with Tukey’s *post hoc* test using R statistical language (R Development Core Team, 2008); different groups are marked with letters at the 0.05 significance level. For all error bars, data are mean ± S.E.M. *P* values were generated using two-tailed Student’s *t*-tests; N/S, *P* ≥ 0.05, \**P* < 0.05, \*\**P* < 0.01, \*\*\**P* < 0.001.

### Accession numbers

The Arabidopsis AGI locus identifiers of genes used in this article are CDC48A (AT3G09840), CDC48B (AT3G53230), CDC48C (AT5G03340), GID1A (AT3G05120), GID1B (AT5G27320), GID1C (AT3G63010), NPL4B (AT3G63000), and UFD1B (AT2G21270)

## Supplementary Figures

**Supplementary Fig. S1. Blue light promotes the nuclear accumulation of CDC48 without influencing its transcript levels a**, Representative images acquired at the end of the experiment of tobacco leaves co-transformed with *35S::turboID-GFP-CDC48A* and the nuclear marker *35S::NLS-Cherry*, grown under continuous blue light. Sum-intensity Z-projections are shown. Scale bars, 50 µm. **b**, Representative images of the live-cell imaging of Col-0 and *cry1-304* seedlings transformed with 35S::YFP-CDC48A, grown under continuous blue light for 4 d. Scale bars, 100 µm. **c**, RNA-seq coverage plot showing transcript levels of *CDC48A* in 6-day-old Col-0 seedlings grown at 22 °C under continuous darkness or red light. Data were reanalysed from a publicly available dataset deposited in the Gene Expression Omnibus (GEO) under the accession number GSE246071. All reads are presented as counts per million (CPM), and genomic coordinates are shown in base pairs (bp).

**Supplementary Fig. S2.** Expression of *^YFP^CDC48A* restores normal hypocotyl elongation in the *cdc48a-4* background, whereas *cdc48b* and *cdc48c* mutants exhibit WT-like growth a, Representative images of *35S::YFP-CDC48A* (*CDC48A*-ox), *cdc48a-4*, and *35S::YFP-CDC48A cdc48a-4* (*CDC48A*-ox + *cdc48a-4*) plants grown under long day conditions at 22 °C for 21 d. **b**, Hypocotyl length of Col-0 wild-type, *35S::YFP-CDC48A* (*CDC48A*-ox, lines 1 and 2), *cdc48a-4*, and *35S::YFP-CDC48A cdc48a-4* (*cdc48a-4 CDC48A*-ox, lines 1 and 5) seedlings grown under blue light. **c**, Quantification of hypocotyl length of Col-0, *cdc48a-4*, *cdc48b-2*, and *cdc48c-1* seedlings grown under continuous darkness, red, or blue light. **d**, Hypocotyl lengths of Col-0 and the *cdc48a-4*, *cdc48b-2*, *cdc48b-3*, *cdc48c-1*, *cdc48c-2*, *cdc48c-3*, *cdc48b-2 cdc48c-1*, *cdc48b-3 cdc48c-2*, and *cdc48a-4 cdc48b-3* mutants grown under continuous blue light in the presence or absence of the CDC48 activity inhibitor CB-5083 (1 µM). DMSO (0.05 % v/v) treatment served as a control. For **b** to **d**, seedlings were grown for 4 d at 22 °C. All data are means (±SEM) of *n* = 3-4 independent biological replicates, each consisting of at least 15 seedlings grown on the same plate. Different letters indicate significant differences among means as determined using one-way ANOVA followed by Tukey’s *post-hoc* test (*P*<0.05).

**Supplementary Fig. S3. CDC48 promotes blue light-mediated hypocotyl growth inhibition through the canonical photomorphogenic pathway a-g**, Seedlings were grown under continuous blue light for 4 d at 22 °C in the presence or absence of the CDC48 activity inhibitor CB-5083 (1 µM). DMSO (0.05 % v/v) treatment served as a control. Quantification of hypocotyl lengths in Col-0 and the *cry1-304* (**a**), *cdc48a-4*, *cry1-17* (**b**), *rdr6-12*, *rdr6-12 CRY1*-ox (**c**), *CRY2*-ox (**d**), *cop1-4*, *cop1-6* (**e**), *hy5-221*, *hy5-215* (**f**), *pif1-2*, *pif3-1*, *pif4-2*, *pif4-101*, *pif5-3*, *pif7-1*, *pif1-1 pif3-3*, *pif3-3 pif4-2*, *pif4-101 pif5-3*, *pif3-3 pif4-2 pif5-3*, *pif4-101 pif5-3 pif7-1*, and *pif1-1 pif3-7 pif4-2 pif5-3* (**g**), mutants at the end of the experiment. For **d**, gray numbers indicate fold changes. All data are means (±SEM) of *n* = 3-4 independent biological replicates, each consisting of at least 15 seedlings grown on the same plate. For **g**, statistical analysis was performed using a two-tailed Student’s *t*-test. Different letters indicate significant differences among means as determined using one-way ANOVA followed by Tukey’s *post-hoc* test (*P*<0.05).

**Supplementary Fig. S4. Mutation of *CDC48A* or *NPL4* results in comparable phenotypes under blue light conditions a**, Immunoblot of GFP from 4-day-old *35S::NPL4B-GFP* seedlings grown under continuous darkness, red, or blue light. GAPC served as loading control. **b**, Quantification of hypocotyl length of Col-0 wild-type, *cdc48a-4*, and *npl4a npl4b* (*npl4ab*) seedlings grown under continuous darkness, red, or blue light for 4 d at 22 °C. **c**, Quantification of hypocotyl lengths in Col-0, *npl4ab*, and *ufd1b ufd1c* (*ufd1bc*) seedlings grown under blue light for 4 d at 22 °C in the presence or absence of the CDC48 activity inhibitor CB-5083 (1 µM). DMSO (0.05 % v/v) treatment served as a control. For **b** and **c**, data are means (±SEM) of *n* = 3 independent biological replicates, each consisting of at least 15 seedlings grown on the same plate. Different letters indicate significant differences among means as determined using one-way ANOVA followed by Tukey’s *post-hoc* test (*P*<0.05).

**Supplementary Fig. S5. CDC48A acts on hypocotyl length regulation under blue light independently of auxin** Seedlings were grown under continuous blue light for 4 d at 22 °C. **a**, Quantification of hypocotyl length of Col-0 wild-type, *cdc48a-4*, and *npl4a npl4b* (*npl4ab*) seedlings grown in the presence or absence of the auxin transport inhibitor N-1-naphthylphthalamic acid (NPA; 0.5 µM). **b**, Hypocotyl length of Col-0 and the *cdc48a-4* mutant grown in the presence or absence of the synthetic auxin picloram (PIC; 0.5 µM). **c-d**, Quantification of hypocotyl lengths in Col-0, *yucca6-1D* (**c**), *axr1-3*, and *axr1-12* (**d**) seedlings grown in the presence or absence of the CDC48 activity inhibitor CB-5083 (1 µM). DMSO (0.05 % v/v) treatment served as a control. **e**, Representative images of GUS staining at the end of the experiment for transformed Col-0 seedlings expressing *GUS* under the control of the artificial *DR5* promoter, grown as in **c**. All data are means (±SEM) of *n* = 3-4 independent biological replicates, each consisting of at least 15 seedlings grown on the same plate. Different letters indicate significant differences among means as determined using one-way ANOVA followed by Tukey’s *post-hoc* test (*P*<0.05).

**Supplementary Fig. S6. CDC48A blue light-mediated suppression of hypocotyl via gibberellin homeostasis** Seedlings were grown under continuous blue light for 4 d at 22 °C. **a-b**, Quantification of hypocotyl length of Col-0 wild-type, *ga3ox1-3 ga3ox2-1* (**a**), and *ga1-3* (**b**) seedlings grown in the presence or absence of the CDC48 activity inhibitor CB-5083 (1 µM). **c**, Levels of the indicated GA metabolites in Col-0 and *cdc48a-4* seedlings grown in the presence or absence of CB-5083 (1 µM), expressed as ng GA/g fresh weight. Red labels highlight GA catabolites. **d**, Representative sum-intensity Z-projections of GFP signal from live-cell imaging of *pRGA::GFP-RGA* seedlings grown as in **a**. Scale bars, 250 µm. **e**, Hypocotyl length of Col-0, *cdc48a-4*, and *global della* seedlings grown as in **a**. For **a-c** and **e**, data are means (±SEM) of *n* = 3-4 independent biological replicates, each consisting of at least 15 seedlings grown on the same plate. For **a**, **d**, and **e**, DMSO (0.05 % v/v) treatment served as a control. Statistical analysis was performed using two-tailed Student’s *t*-test or one-way ANOVA, and letters denote significant differences with a Tukey’s *post hoc* test at *P*<0.05.

**Supplementary Fig. S7. PUX1 does not seem to influence hypocotyl elongation under blue light a**, Schematic representation of the *PUX1* gene structure. Exons are shown in sand, untranslated regions in gray, and introns as a gray line. The triangles indicate T-DNA insertions in the *pux1-3* mutant lines. **b**, Quantification of hypocotyl length of Col-0 wild-type and *pux1-3* seedlings grown under continuous blue light for 4 d at 22 °C. Data are means (±SEM) of *n* = 4 independent biological replicates, each consisting of at least 15 seedlings grown on the same plate. Statistical analysis was performed using two-tailed Student’s *t*-test. ns, not significant.

**Supplementary Fig. S8. Amino-acid alignment of selected UFD1 homologues** Multiple sequence alignment of UFD1 homologues from *Schizosaccharomyces pombe* (SpUFD1, SPBC16A3.09c), *Saccharomyces cerevisiae* (ScUFD1, YGR048W), *Arabidopsis thaliana* (UFD1A, AT2G29070; UFD1B, AT2G21270; UFD1C, AT4G38930), *Oryza sativa* (Os02g08480; Os09g32020; Os01g05120), *Marchantia polymorpha* (Mapoly0015s0157), *Homo sapiens* (HsUFD1, Q92890), *Drosophila melanogaster* (DmUFD1, Q9VTF9), and *Caenorhabditis elegans* (CeUFD-1, Q19584). Conserved residues are shown in white on a red background, with blue boxes indicating conserved amino acid clusters. Residues whose mutations confer thermosensitive phenotypes in fission yeast are highlighted, and the specific residues mutated in AtUFD1B for this study are indicated by blue arrows.

**Supplementary Fig. S9. GID1B does not seem to be expressed in hypocotyls under blue light a**, *GID1B* transcript levels quantified by RT-qPCR in Col-0 wild-type and *cdc48a-4* seedlings grown under continuous blue light for 4 d at 22 °C, in the presence or absence of the CDC48 activity inhibitor CB-5083 (1 µM). DMSO (0.05 % v/v) treatment served as a control. Data were normalized to *ACTIN2/8* transcript levels and are shown on a log_2_ scale relative to Col-0 grown in DMSO, which was set to zero. Data are presented as floating bar plots, with the line indicating the mean of *n* = 3 independent biological replicates and individual values shown. Different letters indicate significant differences among means as determined using one-way ANOVA followed by Tukey’s *post-hoc* test (*P*<0.05). **b**, Representative images of GUS staining for transformed Col-0 seedlings expressing *GID1B* fused to the *GUS* reporter under the control of *GID1B* endogenous promoter (*pGID1B::GID1B-GUS*), grown as in **a**. Scale bars, 500 µm.

## Supplementary data

Supplementary Table S1: PISA analysis of UFD1B-GID1A predicted structure.

Supplementary Table S2: Set of primers used in this study for genotyping.

Supplementary Table S3: Set of primers used in this study for cloning.

Supplementary Table S4: Plasmids used in this study.

Supplementary Table S5: Set of primers used in this study for RT-qPCR experiments.

## Acknowledgments

The authors are grateful to the plant community for their willingness to share seed material with us. This work used computational resources from UNC Supercómputo (CCAD) – Universidad Nacional de Córdoba (https://supercomputo.unc.edu.ar), which are part of SNCAD, Argentina. The authors thank Dr. Elina Welchen for her help with the setting of some experiments and Valentina Correa for her assistance. We thank Dr. Raquel L. Chan for her constant support.

## Author contributions

M.C. conceived the study. A.L.A. and M.C. designed experiments. A.L.Ar. performed the Alphafold protein structure prediction. C.B. obtained single and double mutants and the genetic constructs for *CDC48B* and *CDC48C*. M.D.G. genotyped *gid1* single and double mutants. M.D.G., E.C., and M.A.P.-A. quantified and analyzed gibberellin levels. M.C. contributed to the Y2H and hypocotyl quantification studies. A.L.A. performed all other experiments. A.L.A. and M.C. analyzed the project. M.C. supervised the project. O.L., M.A.P.-A., and M.C. acquired funding. A.L.A. and M.C. conceived and wrote the manuscript. All the authors contributed to the data analysis and discussion and revised the manuscript.

## Data availability

All data supporting the findings of this study, including supplementary materials, are available from the corresponding author upon request.

