## Supplementary Data for "The segregase CDC48 integrates blue light and hormonal cues to regulate photomorphogenesis in Arabidopsis"

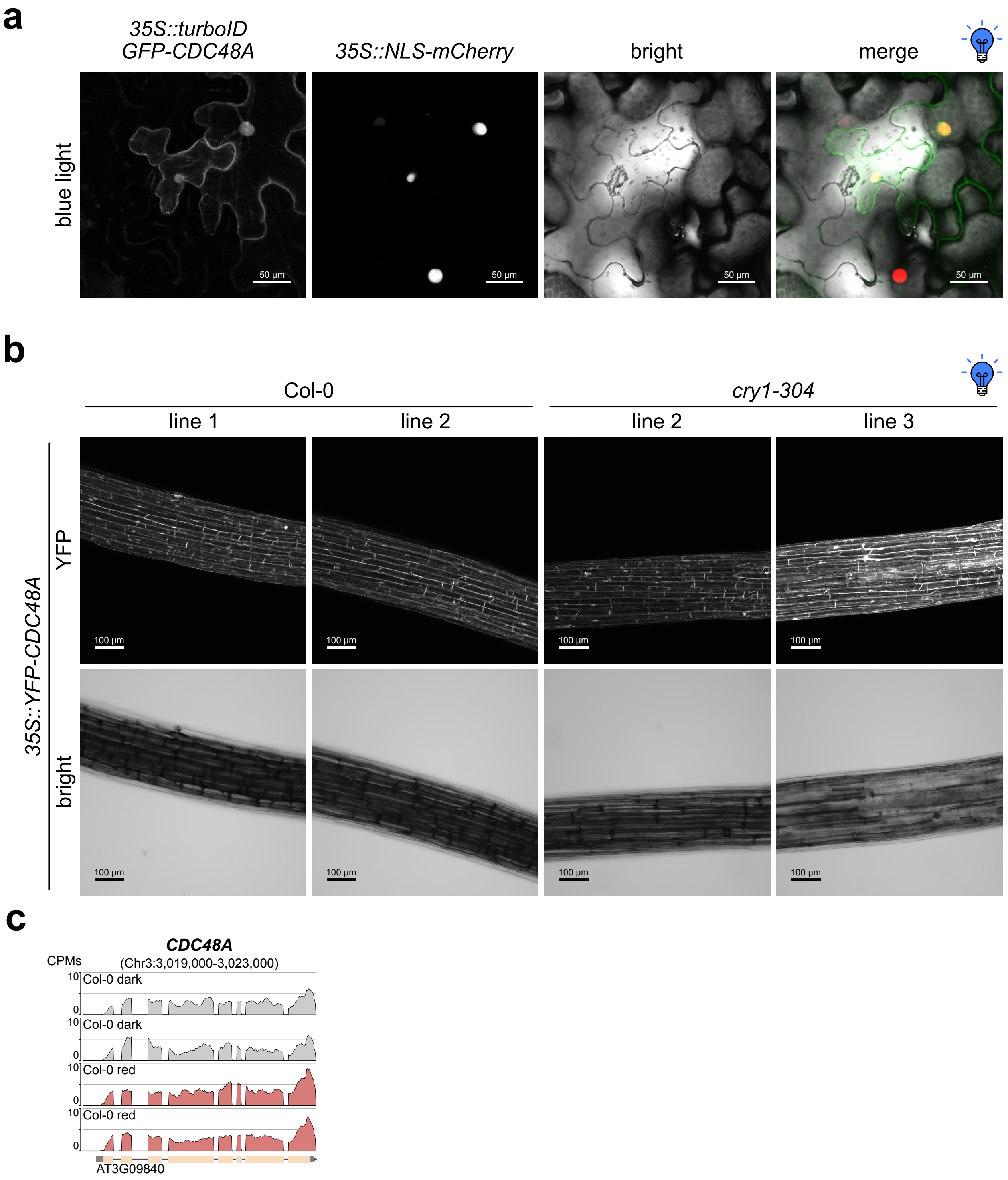


**Supplementary Fig. S1. Blue light promotes the nuclear accumulation of CDC48 without influencing its transcript levels**

**a**, Representative images acquired at the end of the experiment of tobacco leaves co-transformed with *35S::turboID-GFP-CDC48A* and the nuclear marker *35S::NLS-Cherry*, grown under continuous blue light. Sum-intensity Z-projections are shown. Scale bars, 50 µm. **b**, Representative images of the live-cell imaging of Col-0 and *cry1-304* seedlings transformed with *35S::YFP-CDC48A*, grown under continuous blue light for 4 d. Scale bars, 100 µm. **c**, RNA-seq coverage plot showing transcript levels of *CDC48A* in 6-day-old Col-0 seedlings grown at 22 ˚C under continuous darkness or red light. Data were reanalysed from a publicly available dataset deposited in the Gene Expression Omnibus (GEO) under the accession number GSE246071. All reads are presented as counts per million (CPM), and genomic coordinates are shown in base pairs (bp).

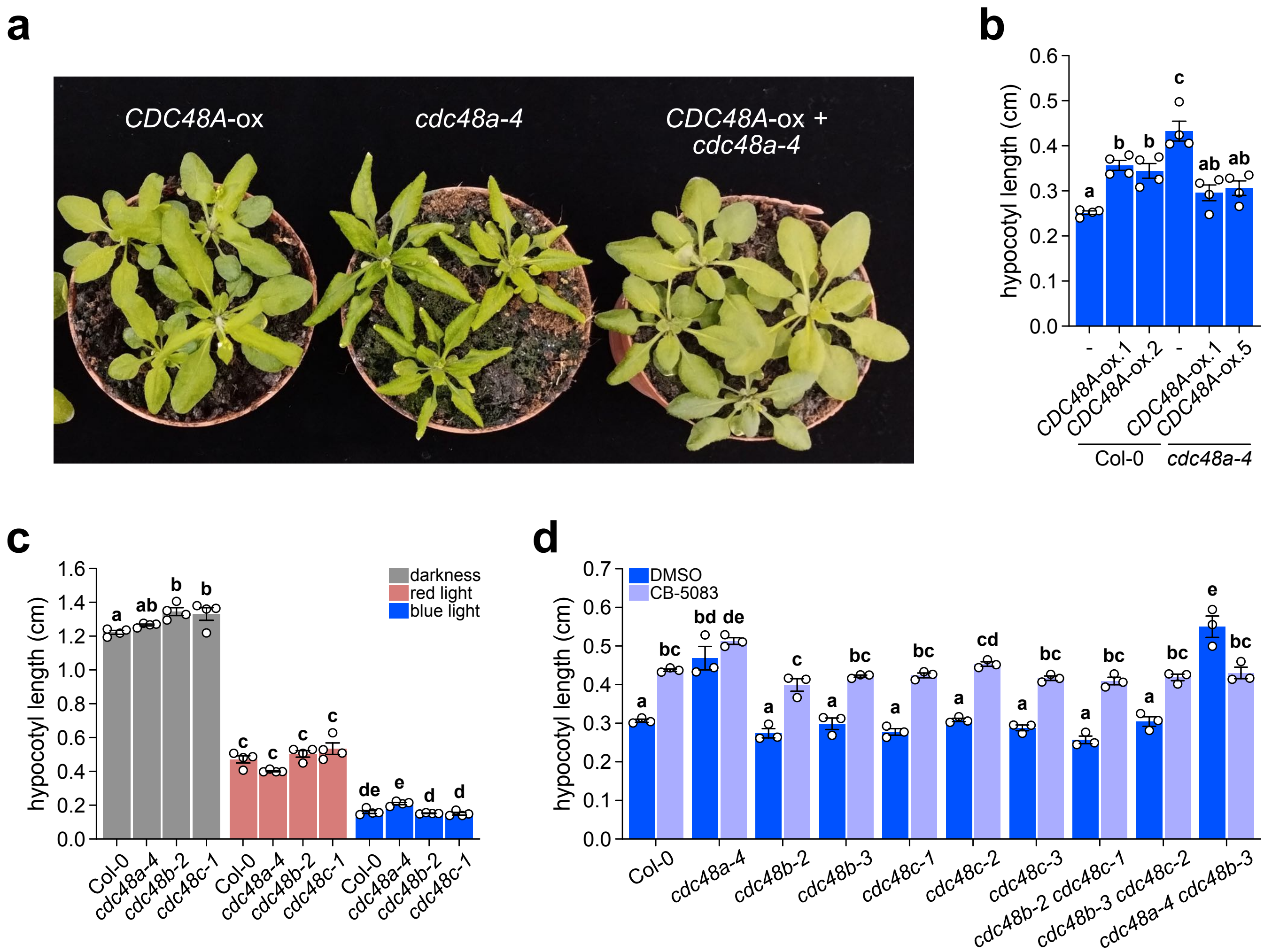


**Supplementary Fig. S2. Expression of *$^{YFP}$CDC48A* restores normal hypocotyl elongation in the *cdc48a-4* background, whereas *cdc48b* and *cdc48c* mutants exhibit WT-like growth**

**a**, Representative images of *35S::YFP-CDC48A* (*CDC48A*-ox), *cdc48a-4*, and *35S::YFP-CDC48A cdc48a-4* (*CDC48A*-ox + *cdc48a-4*) plants grown under long day conditions at 22 ˚C for 21 d. **b**, Hypocotyl length of Col-0 wild-type, *35S::YFP-CDC48A* (*CDC48A*-ox, lines 1 and 2), *cdc48a-4*, and *35S::YFP-CDC48A cdc48a-4* (*cdc48a-4 CDC48A*-ox, lines 1 and 5) seedlings grown under blue light. **c**, Quantification of hypocotyl length of Col-0, *cdc48a-4*, *cdc48b-2*, and *cdc48c-1* seedlings grown under continuous darkness, red, or blue light. **d**, Hypocotyl lengths of Col-0 and the *cdc48a-4*, *cdc48b-2*, *cdc48b-3*, *cdc48c-1*, *cdc48c-2*, *cdc48c-3*, *cdc48b-2 cdc48c-1*, *cdc48b-3 cdc48c-2*, and *cdc48a-4 cdc48b-3* mutants grown under continuous blue light in the presence or absence of the CDC48 activity inhibitor CB-5083 (1 µM). DMSO (0.05 % v/v) treatment served as a control. For **b** to **d**, seedlings were grown for 4 d at 22 ˚C. All data are means (±SEM) of *n* = 3-4 independent biological replicates, each consisting of at least 15 seedlings grown on the same plate. Different letters indicate significant differences among means as determined using one-way ANOVA followed by Tukey's *post-hoc* test ($P$<0.05).

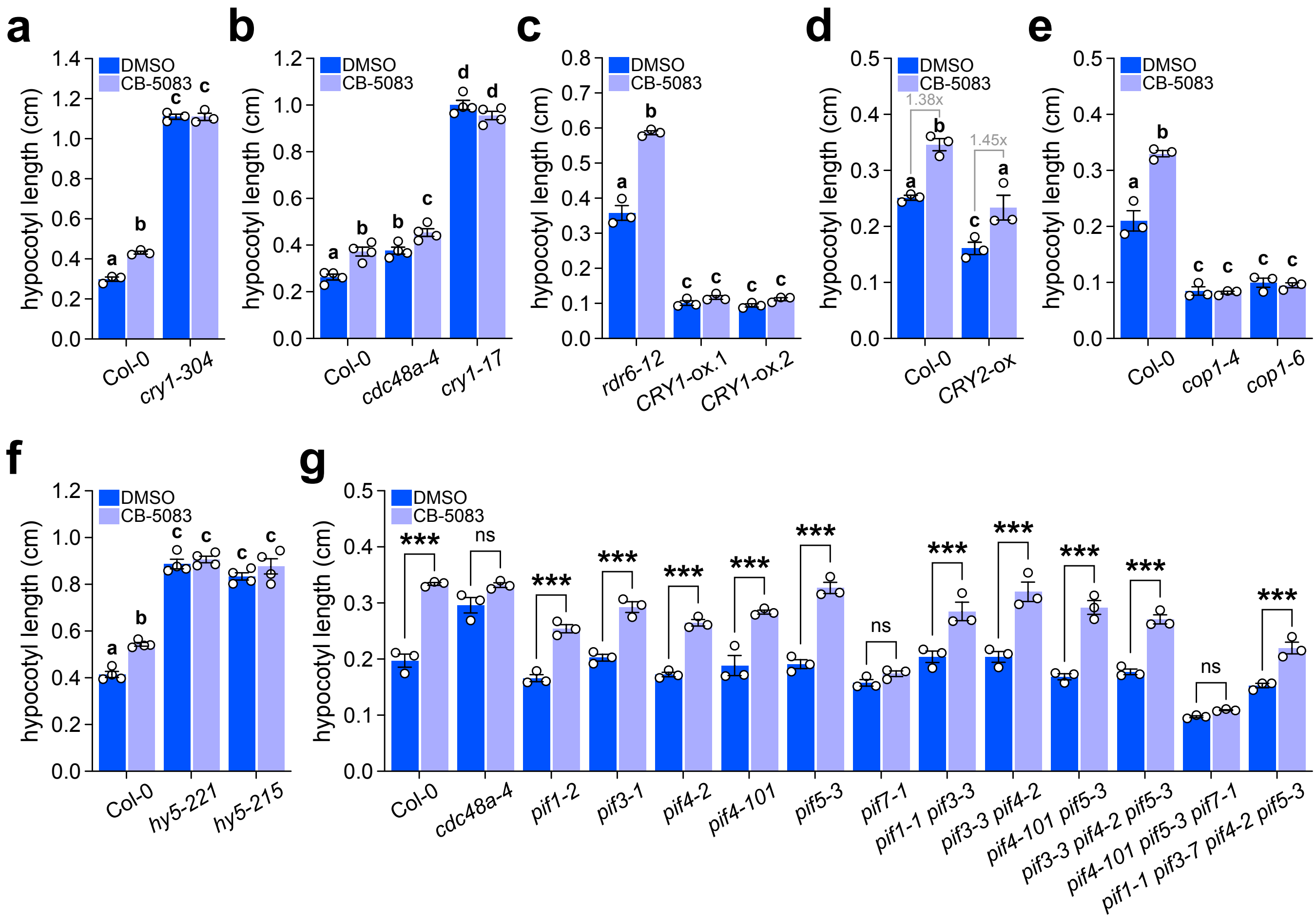


**Supplementary Fig. S3. CDC48 promotes blue light-mediated hypocotyl growth inhibition through the canonical photomorphogenic pathway**

**a-g**, Seedlings were grown under continuous blue light for 4 d at 22 ˚C in the presence or absence of the CDC48 activity inhibitor CB-5083 (1 µM). DMSO (0.05 % v/v) treatment served as a control. Quantification of hypocotyl lengths in Col-0 and the *cry1-304* (**a**), *cdc48a-4*, *cry1-17* (**b**), *rdr6-12*, *rdr6-12 CRY1*-ox (**c**), *CRY2*-ox (**d**), *cop1-4*, *cop1-6* (**e**), *hy5-221*, *hy5-215* (**f**), *pif1-2*, *pif3-1*, *pif4-2*, *pif4-101*, *pif5-3*, *pif7-1*, *pif1-1 pif3-3*, *pif3-3 pif4-2*, *pif4-101 pif5-3*, *pif3-3 pif4-2 pif5-3*, *pif4-101 pif5-3 pif7-1*, and *pif1-1 pif3-7 pif4-2 pif5-3* (**g**), mutants at the end of the experiment. For **d**, gray numbers indicate fold changes. All data are means (±SEM) of $n$ = 3-4 independent biological replicates, each consisting of at least 15 seedlings grown on the same plate. For **g**, statistical analysis was performed using a two-tailed Student's $t$-test. Different letters indicate significant differences among means as determined using one-way ANOVA followed by Tukey's *post-hoc* test ($P$<0.05).

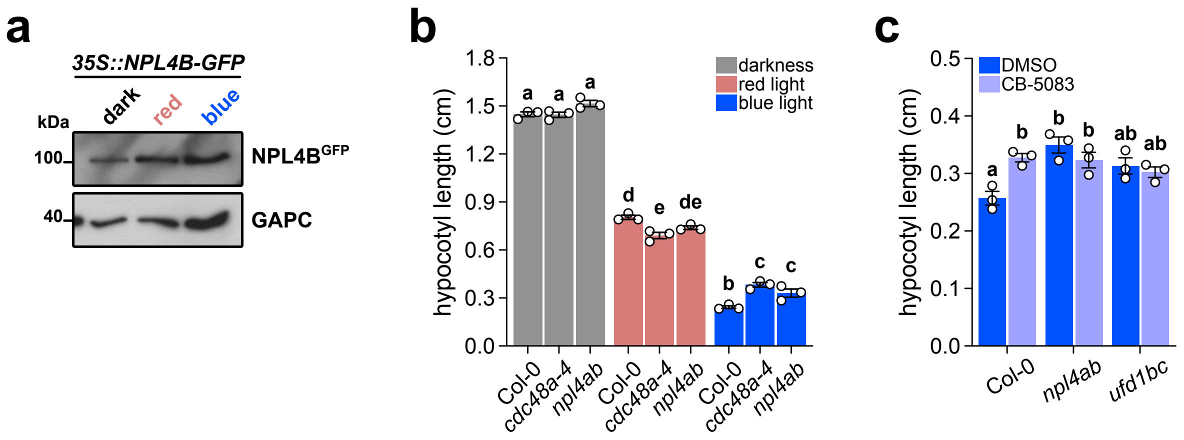


**Supplementary Fig. S4. Mutation of *CDC48A* or *NPL4* results in comparable phenotypes under blue light conditions**
**a**, Immunoblot of GFP from 4-day-old *35S::NPL4B-GFP* seedlings grown under continuous darkness, red, or blue light. GAPC served as loading control. **b**, Quantification of hypocotyl length of Col-0 wild-type, *cdc48a-4*, and *npl4a npl4b* (*npl4ab*) seedlings grown under continuous darkness, red, or blue light for 4 d at 22 ˚C. **c**, Quantification of hypocotyl lengths in Col-0, *npl4ab*, and *ufd1b ufd1c* (*ufd1bc*) seedlings grown under blue light for 4 d at 22 ˚C in the presence or absence of the CDC48 activity inhibitor CB-5083 (1 µM). DMSO (0.05 % v/v) treatment served as a control. For **b** and **c**, data are means (±SEM) of *n* = 3 independent biological replicates, each consisting of at least 15 seedlings grown on the same plate. Different letters indicate significant differences among means as determined using one-way ANOVA followed by Tukey's *post-hoc* test (*P*<0.05).

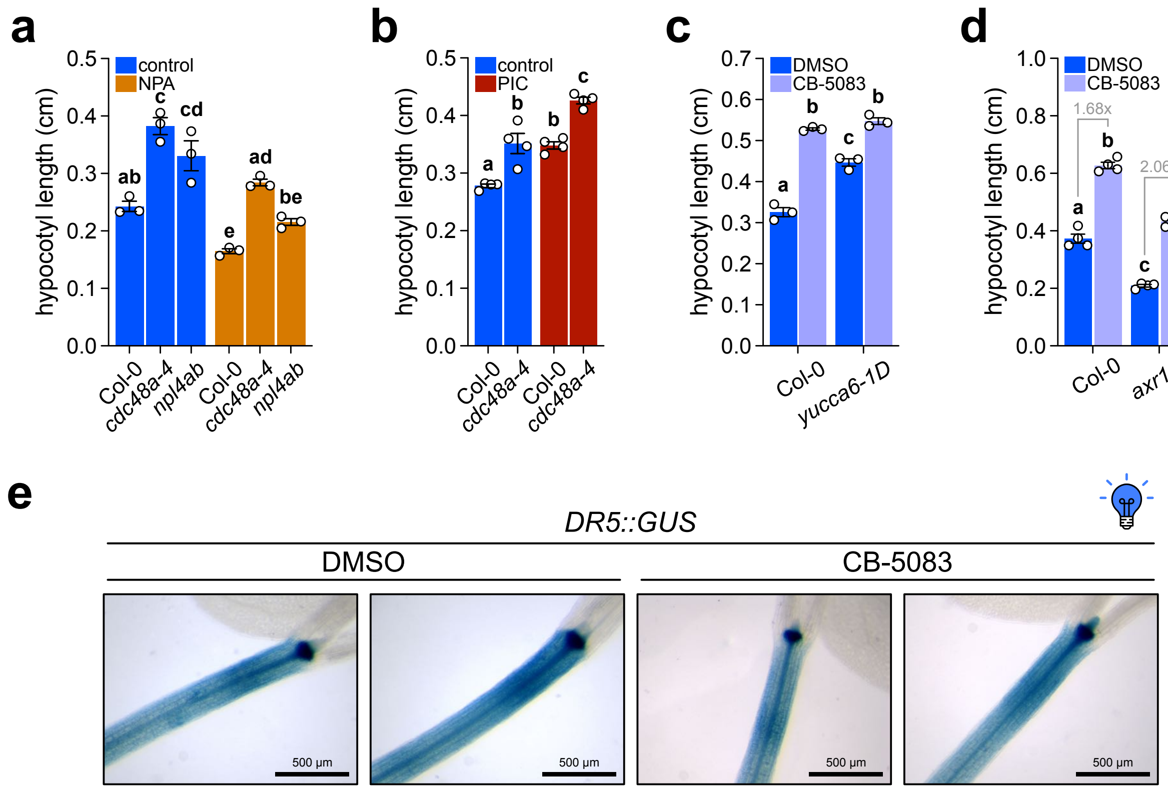


**Supplementary Fig. S5. CDC48A acts on hypocotyl length regulation under blue light independently of auxin**
Seedlings were grown under continuous blue light for 4 d at 22 ˚C. **a**, Quantification of hypocotyl length of Col-0 wild-type, *cdc48a-4*, and *npl4a npl4b* (*npl4ab*) seedlings grown in the presence or absence of the auxin transport inhibitor N-1-naphthylphthalamic acid (NPA; 0.5 µM). **b**, Hypocotyl length of Col-0 and the *cdc48a-4* mutant grown in the presence or absence of the synthetic auxin picloram (PIC; 0.5 µM). **c-d**, Quantification of hypocotyl lengths in Col-0, *yucca6-1D* (**c**), *axr1-3*, and *axr1-12* (**d**) seedlings grown in the presence or absence of the CDC48 activity inhibitor CB-5083 (1 µM). DMSO (0.05 % v/v) treatment served as a control. **e**, Representative images of GUS staining at the end of the experiment for transformed Col-0 seedlings expressing *GUS* under the control of the artificial *DR5* promoter, grown as in **c**. All data are means (±SEM) of $n$ = 3-4 independent biological replicates, each consisting of at least 15 seedlings grown on the same plate. Different letters indicate significant differences among means as determined using one-way ANOVA followed by Tukey's *post-hoc* test ($P$<0.05).

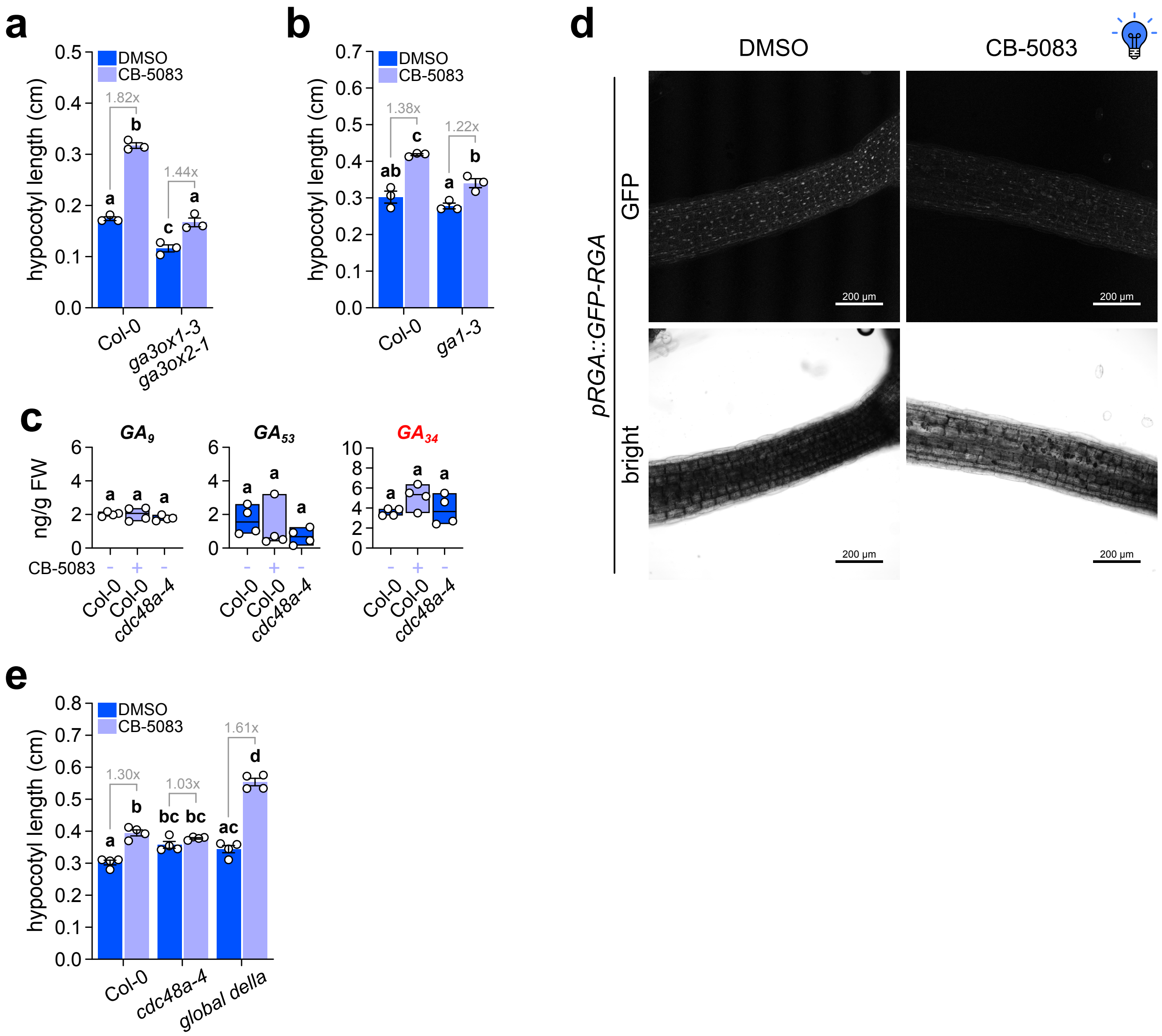


**Supplementary Fig. S6. CDC48A blue light-mediated suppression of hypocotyl via gibberellin homeostasis**
Seedlings were grown under continuous blue light for 4 d at 22 ˚C. **a-b**, Quantification of hypocotyl length of Col-0 wild-type, *ga3ox1-3 ga3ox2-1* (**a**), and *ga1-3* (**b**) seedlings grown in the presence or absence of the CDC48 activity inhibitor CB-5083 (1 µM). **c**, Levels of the indicated GA metabolites in Col-0 and *cdc48a-4* seedlings grown in the presence or absence of CB-5083 (1 µM), expressed as ng GA/g fresh weight. Red labels highlight GA catabolites. **d**, Representative sum-intensity Z-projections of GFP signal from live-cell imaging of *pRGA::GFP-RGA* seedlings grown as in **a**. Scale bars, 250 µm. **e**, Hypocotyl length of Col-0, *cdc48a-4*, and *global della* seedlings grown as in **a**. For **a-c** and **e**, data are means (±SEM) of $n$ = 3-4 independent biological replicates, each consisting of at least 15 seedlings grown on the same plate. For **a**, **d**, and **e**, DMSO (0.05 % v/v) treatment served as a control. Statistical analysis was performed using two-tailed Student's $t$-test or one-way ANOVA, and letters denote significant differences with a Tukey's *post hoc* test at $P$<0.05.

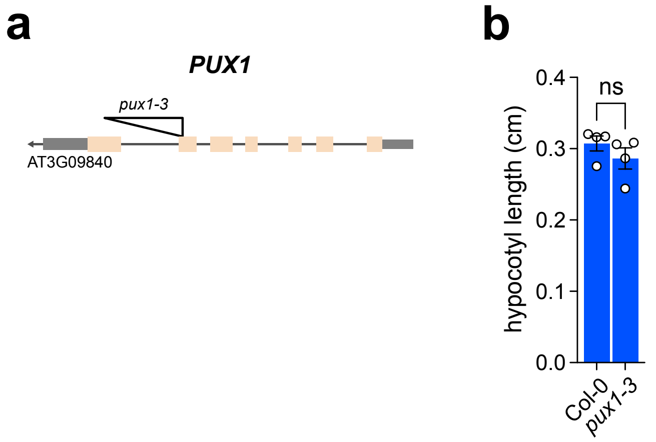


**Supplementary Fig. S7. PUX1 does not seem to influence hypocotyl elongation under blue light**
**a**, Schematic representation of the *PUX1* gene structure. Exons are shown in sand, untranslated regions in gray, and introns as a gray line. The triangles indicate T-DNA insertions in the *pux1-3* mutant lines. **b**, Quantification of hypocotyl length of Col-0 wild-type and *pux1-3* seedlings grown under continuous blue light for 4 d at 22 ˚C. Data are means (±SEM) of $n$ = 4 independent biological replicates, each consisting of at least 15 seedlings grown on the same plate. Statistical analysis was performed using two-tailed Student's *t*-test. ns, not significant.

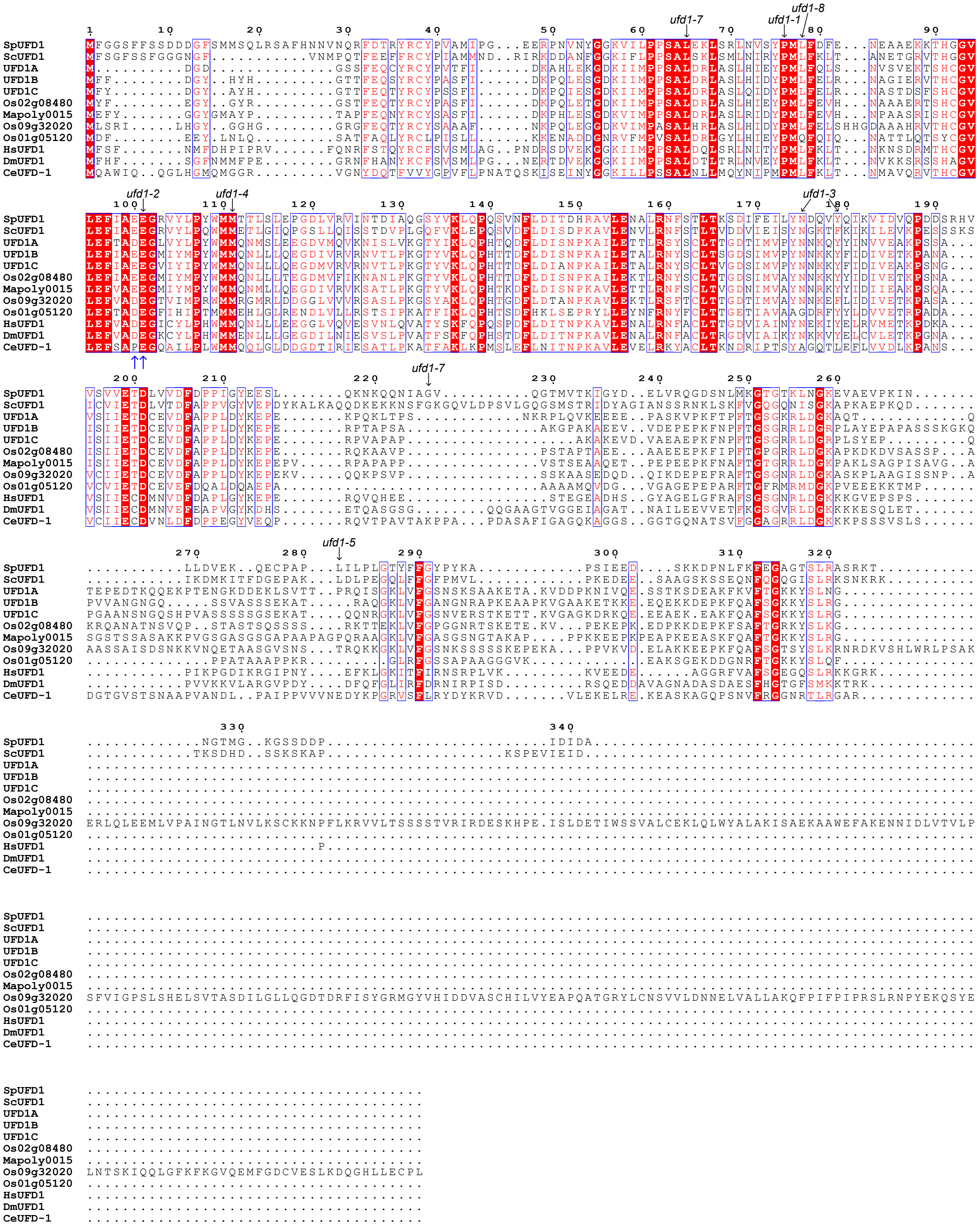


**Supplementary Fig. S8. Amino-acid alignment of selected UFD1 homologues**

Multiple sequence alignment of UFD1 homologues from *Schizosaccharomyces pombe* (SpUFD1, SPBC16A3.09c), *Saccharomyces cerevisiae* (ScUFD1, YGR048W), *Arabidopsis thaliana* (UFD1A, AT2G29070; UFD1B, AT2G21270; UFD1C, AT4G38930), *Oryza sativa* (Os02g08480; Os09g32020; Os01g05120), *Marchantia polymorpha* (Mapoly0015s0157), *Homo sapiens* (HsUFD1, Q92890), *Drosophila melanogaster* (DmUFD1, Q9VTF9), and *Caenorhabditis elegans* (CeUFD-1, Q19584). Conserved residues are shown in white on a red background, with blue boxes indicating conserved amino acid clusters. Residues whose mutations confer thermosensitive phenotypes in fission yeast are highlighted, and the specific residues mutated in AtUFD1B for this study are indicated by blue arrows.

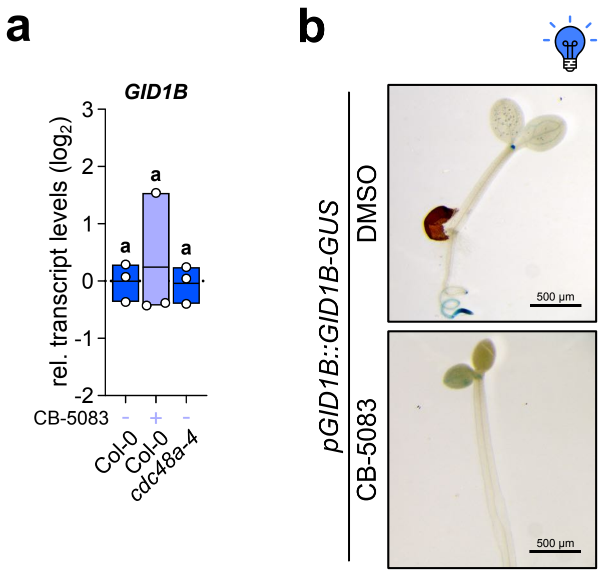


**Supplementary Fig. S9. GID1B does not seem to be expressed in hypocotyls under blue light**

**a**, *GID1B* transcript levels quantified by RT-qPCR in Col-0 wild-type and *cdc48a-4* seedlings grown under continuous blue light for 4 d at 22 ˚C, in the presence or absence of the CDC48 activity inhibitor CB-5083 (1 µM). DMSO (0.05 % v/v) treatment served as a control. Data were normalized to *ACTIN2/8* transcript levels and are shown on a $\log_2$ scale relative to Col-0 grown in DMSO, which was set to zero. Data are presented as floating bar plots, with the line indicating the mean of $n$ = 3 independent biological replicates and individual values shown. Different letters indicate significant differences among means as determined using one-way ANOVA followed by Tukey's *post-hoc* test ($P$<0.05). **b**, Representative images of GUS staining for transformed Col-0 seedlings expressing *GID1B* fused to the *GUS* reporter under the control of *GID1B* endogenous promoter (*pGID1B::GID1B-GUS*), grown as in **a**. Scale bars, 500 µm.

**Supplementary Table S2.** Set of primers used in this study for genotyping

| Locus | Name | Sequence |
|---|---|---|
| - | SALK LBb1.3 | 5'-ATTTTGCCGATTTCGGAAC-3' |
| - | LB3 T-DNA SAIL | 5'-TAGCATCTGAATTTCATAACCAATCTCGATACAC-3' |
| - | GABI-LB-o8474 | 5'-ATAATAACGCTGCGGACATCTACATTTT-3' |
| AT3G09840 | CDC48A-WT F | 5'-CCTTTTTCTTCTGTATCAACGG-3' |
| AT3G09840 | CDC48A-mut F | 5'-CCTTTTTCTTCTGTATCAACGA-3' |
| AT3G09840 | CDC48A-WT R | 5'-CCTGGAAAACAACCACAAGAAG-3' |
| AT3G53230 | CDC48B SAIL_566E10_FP | 5'-GAGATCATCATCATCCGCTG-3' |
| AT3G53230 | CDC48B SAIL_566E10_RP | 5'-GAGATATTCGACAAGGCACG-3' |
| AT3G53230 | CDC48B GABI_FP | 5'-TTCTCATCCCTCCTCTAACTAGGA-3' |
| AT3G53230 | CDC48B GABI_RP | 5'-TAAATATGAACACCGAGCTCCTCA-3' |
| AT5G03340 | CDC48C SAIL_1182E09_FP | 5'-CGAGGCGATACAATTCTCAT-3' |
| AT5G03340 | CDC48C SAIL_1182E09_RP | 5'-CCAATTCCCTAATCTGAGCC-3' |
| AT5G03340 | CDC48C SALK_FP | 5'-GGTTGCTCTCACTCTCACCAG-3' |
| AT5G03340 | CDC48C SALK_RP | 5'-TATGTCAAACGAACCGGAATC-3' |
| AT3G27310 | PUX1 SAILseq_595_F08.1 LP | 5'-TTTTTACCGCCTTTTGGCTAC-3' |
| AT3G27310 | PUX1 SAILseq_595_F08.1 RP | 5'-CACTAGAGTCAACCGTGGCTG-3' |

**Supplementary Table S3.** Set of primers used in this study for cloning.

| Locus | Name | Sequence | Restriction site | Construct |
|---|---|---|---|---|
| AT3G09840 | CDC48A F | 5'-GCGCGGTACCATGTCTACCCCAGCTGAATCTTC-3' | *Kpn*I | pENTR |
| AT3G09840 | CDC48A NdeI F | 5'-GCGCCATATGTCTACCCCAGCTGAATC-3' | *Nde*I | Y2H |
| AT3G09840 | CDC48A R | 5'-GCGCGTCGACACTAATTGTAGAGATCATCATCG-3' | *Sal*I | pENTR and Y2H |
| AT3G09840 | CDC48A G274E F | 5'-TGTATCAACG**A**ACCTGAGATCATGTCCAAATTG-3' | - | mutation |
| AT3G09840 | CDC48A G274E R | 5'-GATCTCAGGT**T**CGTTGATACAGAAGAAAAAGGC-3' | - | mutation |
| AT3G09840 | CDC48A E581Q F | 5'-TTTCTTTGAT**C**AGCTCGACTCCATTGCAACTC-3' | - | mutation |
| AT3G09840 | CDC48A E581Q R | 5'-GAGTCGAGCT**G**ATCAAAGAAAAGAACACATGG-3' | - | mutation |
| AT4G08920 | CRY1 EcoRI F | 5'-GCGCGAATTCGAGATGTCTGGTTCTGTATCTG-3' | *Eco*RI | pENTR |
| AT4G08920 | CRY1 XhoI R | 5'-GCGCCTCGAGTTACCCGGTTTGTGAAAGCCG-3' | *Xho*I | pENTR |
| AT3G05120 | GID1A F | 5'-GCGGAATTCATGGCTGCGAGCGATGAAGTT-3' | *Eco*RI | pENTR and Y2H |
| AT3G05120 | GID1A R | 5'-GCGGTCGACCCAGTGTTAACATTCCGCGT-3' | *Sal*I | pENTR and Y2H |
| AT3G05120 | GID1A woSTOP R | 5'-GCGGTCGACACATTCCGCGTTTACAAACGCC-3' | *Sal*I | pENTR |
| AT5G27320 | GID1C F | 5'-GCGGAATTCTCCATGGCTGGAAGTGAAGAAG-3' | *Eco*RI | Y2H |
| AT5G27320 | GID1C R | 5'-GCGGTCGACGTGTTCTCATTGGCATTCTGCG-3' | *Sal*I | Y2H |
| AT3G63000 | NPL4B F | 5'-GCGCGAATTCATGACGATGCTCAGAGTCCG-3' | EcoRI | pENTR and Y2H |
| AT3G63000 | NPL4B R | 5'-GCGCCTCGAGTTAAGAAGTATTGGCCATGGAG-3' | XhoI | pENTR and Y2H |
| AT3G63000 | NPL4B woSTOP R | 5'-GCGCTCGAGAGAAGTATTGGCCATGGAGTCG-3' | XhoI | pENTR |
| AT2G21270 | UFD1B F | 5'-GCGCGAATTCATGTTTTTCGATGGATACCATTATC-3' | EcoRI | pENTR and Y2H |
| AT2G21270 | UFD1B R | 5'-GCGCCTCGAGTTGATCAACCCCTCAATGAATAC-3' | XhoI | pENTR and Y2H |
| AT2G21270 | UFD1B woSTOP R | 5'-GCGCTCGAGACCCCTCAATGAATACTTCTTCC-3' | XhoI | pENTR |
| AT2G21270 | UFD1B E78A E79A F | 5'-TCATTGCAG**C**AG**C**AGGCATGATTTACATGCCATA-3' | - | mutation |
| AT2G21270 | UFD1B E78A E79A R | 5'-ATCATGCCT**G**CT**G**CTGCAATGAACTCAAGGACTC-3' | - | mutation |

**Supplementary Table S4.** Plasmids used in this study.

| Construct | Source |
|---|---|
| pGADT7 | Clontech® |
| pGADT7 CDC48A | This study |
| pGADT7 GID1A | This study |
| pGADT7 GID1C | This study |
| pGADT7 RGA | |
| pGBKT7 | Clontech® |
| pGBKT7 NPL4B | This study |
| pGBKT7 UFD1B | This study |
| pGBKT7 UFD1B E78A E79A | This study |
| pFAST-R06 | (Shimada *et al.*, 2010) |
| pFAST-R06 35S::GFP-CRY1 | This study |
| pJV-117 | (Hellens *et al.*, 2000) |
| pJV-117 35S::mCherry-GID1A | This study |
| pGWB421 | (Nakagawa *et al.*, 2007); Addgene #74815 |
| pGWB421 35S::Myc-GID1A | This study |
| pGW-GFP-TurboID-GOI | (Tan *et al.*, 2024); Addgene #209381 |
| pGW-GFP-TurboID-GOI 35S::GFP-turboID-CDC48A | This study |
| pGW-GFP-TurboID-GOI 35S::GFP-turboID-CDC48A E581Q | This study |
| pGW-GFP-TurboID-GOI 35S::GFP-turboID-UFD1B | This study |
| pB7WGY2 | (Karimi *et al.*, 2005) |
| pB7WGY2 35S::YFP-CDC48A | This study |
| pAS-054 | (Hellens *et al.*, 2000) |
| pAS-054 35S::NPL4B-N-mCitrine | This study |
| pAS-054 35S::PIF4-N-mCitrine | (Ferrero *et al.*, 2019) |
| pAS-059 | (Hellens *et al.*, 2000) |
| pAS-059 35S::GID1A-C-mCitrine | This study |
| pAS-059 35S::PIF4-C-mCitrine | (Ferrero *et al.*, 2019) |

**Supplementary Table S5.** Set of primers used in this study for RT-qPCR experiments.

| Locus | Name | Sequence |
|---|---|---|
| AT3G18780/AT1G49240 | ACTIN2/8 qF | 5'-GGTAACATTGTGCTCAGTGGTGG-3' |
| AT3G18780/AT1G49240 | ACTIN2/8 qR | 5'-AACGACCTTAATCTTCATGCTGC-3' |
| AT3G09840 | CDC48A qF | 5'-GGAGAAGAGGAGGAGCGAGA-3' |
| AT3G09840 | CDC48A qR | 5'-ACACTCCTACGCGCATACTT-3' |
| AT1G15550 | GA3OX1 qF | 5'-CCCAACATCACCTCAACTACTGC-3' |
| AT1G15550 | GA3OX1 qR | 5'-ACTTGTAGACCGGCGGTATTGTT-3' |
| AT4G25420 | GA20OX1 qF | 5'-AGGACGCTGCTGGACTTCAC-3' |
| AT4G25420 | GA20OX1 qR | 5'-GCCATTGCTACTGGTGCTGAG-3' |
| AT1G78440 | GA2OX1 qF | 5'-CGTTTCCGCAGAGCTAGTCTCTG-3' |
| AT1G78440 | GA2OX1 qR | 5'-AGTGGACCCGAACCGGAATCATG-3' |
| AT3G05120 | GID1A qF | 5'-GAAATGGCTGCGAGCGATGAAG-3' |
| AT3G05120 | GID1A qR | 5'-ATTGAGAGGAACCACTGTTCTGC-3' |
| AT3G63010 | GID1B qF | 5'-TGGAGACTATGGCTGGTGGTAAC-3' |
| AT3G63010 | GID1B qR | 5'-AGTGGGACAATTCTCTTGCATTCG-3' |
| AT5G27320 | GID1C qF | 5'-TCTTCGATCTGGGCTTTCGTGTC-3' |
| AT5G27320 | GID1C qR | 5'-ATTGAGAGGAACCACTGTCTTGC-3' |
